# Integrated tumour-immune cell response modelling of Luminal A breast cancer details malignant signalling and ST3Gal1 inhibitor-induced reversal

**DOI:** 10.1101/2025.04.03.647001

**Authors:** Hikmet Emre Kaya, Kevin J. Naidoo

## Abstract

Aberrant O-glycosylation of mucin-type glycopeptide 1 (MUC1) is implicated in cancerous cellular processes involving the manipulation of immune response to favour tumour growth and metastasis. There is an unmet need for systems glycobiology models to probe the relationship between MUC1 O-glycosylation and immune cells within the tumour microenvironment. We expand on the sparsely understood MUC1 and immune cell interactions by building a complete systems model that combines the glycosylation network in the tumour cell with downstream biological networks. An ordinary differential equations-based model of the effect of aberrant glycosylation on immune modulation in breast cancer was constructed. The model comprises three interdependent component models that are MUC1-type O-glycosylation in the tumour cell, chemokine secretion in macrophages, and signal transduction in the tumour cells. A comparative CytoCopasi algorithm was developed to sequentially perturb the networks by an aberrant O-glycosylation. Comparative simulations revealed that upregulation of tumour-associated MUC1 sialyl-T antigen in Luminal A breast cancer stimulated the upregulation of the chemokine CXCL5 in tumour-associated macrophages. Consequently, increased CXCL5 binding by the tumour cell led to a positive feedback loop through overactive signal transduction and autocrine CXCL5 production. Finally, perturbing the glycosylation network with the sialyltransferase inhibitor Soyasaponin-I abrogated the cancerous upregulations in the downstream networks.

## Introduction

Epithelial cells play a central role in fending off disease introduced by pathogens as the first line of defence against microorganism invasion. Glycoconjugates such as mucins are a primary component of this defence mechanism (Boll et al. 2017; Lindén et al. 2009). In epithelial cancers, the biochemical and molecular defence networks against diseases are altered in these cells to facilitate their transition toward tumour cells. Of particular relevance are the alterations in glycoconjugates where the resultant aberrant glycosylation(s) is significantly associated with poor prognosis and malignancy in cancer (Bangarh et al. 2023; Meany and Chan 2011).

Sialyltransferases such as β-Galactoside α-2,3-Sialyltransferase 1 (ST3Gal1) have been identified as possible targets in cancer care (Burchell et al. 1999). On the other hand, glycosyltransferase inhibitor discovery has been hampered by a lack of structural and mechanistic data to guide the design process(Bowles and Gloster 2021; Gloster and Vocadlo 2012). At the molecular scale, computational work schemes are developed to address this problem(Crous and Naidoo 2016; Senapathi et al. 2019; Senapathi et al. 2020). Alongside this, a small number of potential inhibitors have been derived from soyasaponin I or triazole against the cancer-associated ST3Gal1(Chang et al. 2006; Szabo et al. 2024; Wu et al. 2001). Soyasaponin I- and lithocholic acid-derived inhibitors impeded invasion and migration capabilities in breast cancer cell lines (Chen et al. 2011; Hsu et al. 2005). There is the caveat that these studies involved endpoint assays with substrate concentrations below saturation, yielding unrealistic IC_50_ values. A universal glycosyltransferase assay developed by Nashed et al. uncovered a time-dependent inhibition mechanism for Soyasaponin I (Nashed and Naidoo 2024), revealing that the original study(Wu et al. 2001) overestimated the binding affinity of the inhibitor. Despite the emergence of accurate kinetics assays, the potential role of sialyltransferase inhibitors in mitigating tumourigenic pathways and phenotypes has not been unveiled extensively.

Chemical systems biology models that interface glycosylation with downstream pathways enable a means to interrogate the role of aberrant glycosylation in tumorigenesis and predict the efficacy of sialyltransferase inhibitors at a molecular length scale. Networks of chemical reactions, epitomised by ordinary differential equation (ODE) formalisms, allow for dynamic simulation exemplified by software packages, such as Complex Pathway Simulator (COPASI) (Hoops et al. 2006). More recently, the modelling and simulation tools of COPASI were integrated into Cytoscape (Shannon et al. 2003) through the Java-based application CytoCopasi (Kaya and Naidoo 2023) to make possible comparative network simulation and visualization. Nevertheless, the closed structure of systems glycobiology models remains an impediment to quantifying the effect of glycan alterations on disease progression. Specifically, many ODE models were aimed at isolating glycosylation networks under steady-state assumptions to correlate enzyme activities to glycan distribution, and to optimize a specific glycan epitope for recombinant biotherapeutics production (Kouka et al. 2022; Krambeck and Betenbaugh 2005; Umaña and Bailey 1997). A human physiology-related study conducted by Liu et al. demonstrated the impact of glycosyltransferase activity on sialyl Lewis-X (sLe^x^) formation on P-selectin Glycoprotein Ligand-1 (PSGL-1), a leukocyte surface glycoprotein (Liu et al. 2008). However, the model did not capture the impact of this epitope on leukocyte adhesion and tissue inflammation. To date, no ODE model has been developed to investigate the downstream effects of aberrant glycosylation on a biological organism, a disease and, of relevance here, specifically cancer.

Open models of glycosylation can be established for glycoproteins known to interact with a myriad of intracellular and intercellular biological networks. Among these glycoproteins, Mucin-1 (MUC1), a transmembrane heterodimer glycoprotein, stands out for its overexpression and aberrant glycosylation in cancer, occasioning dysregulated interactions with signalling kinases and the tumour microenvironment (TME). In particular, its N-subunit comprises 20-amino acid-long tandem repeats (TR), which shift from branched core 2-type O-glycans to shorter and sialic acid-bearing core 1-type O-glycans in cancer (Fig. 1).

**Fig. 1:**
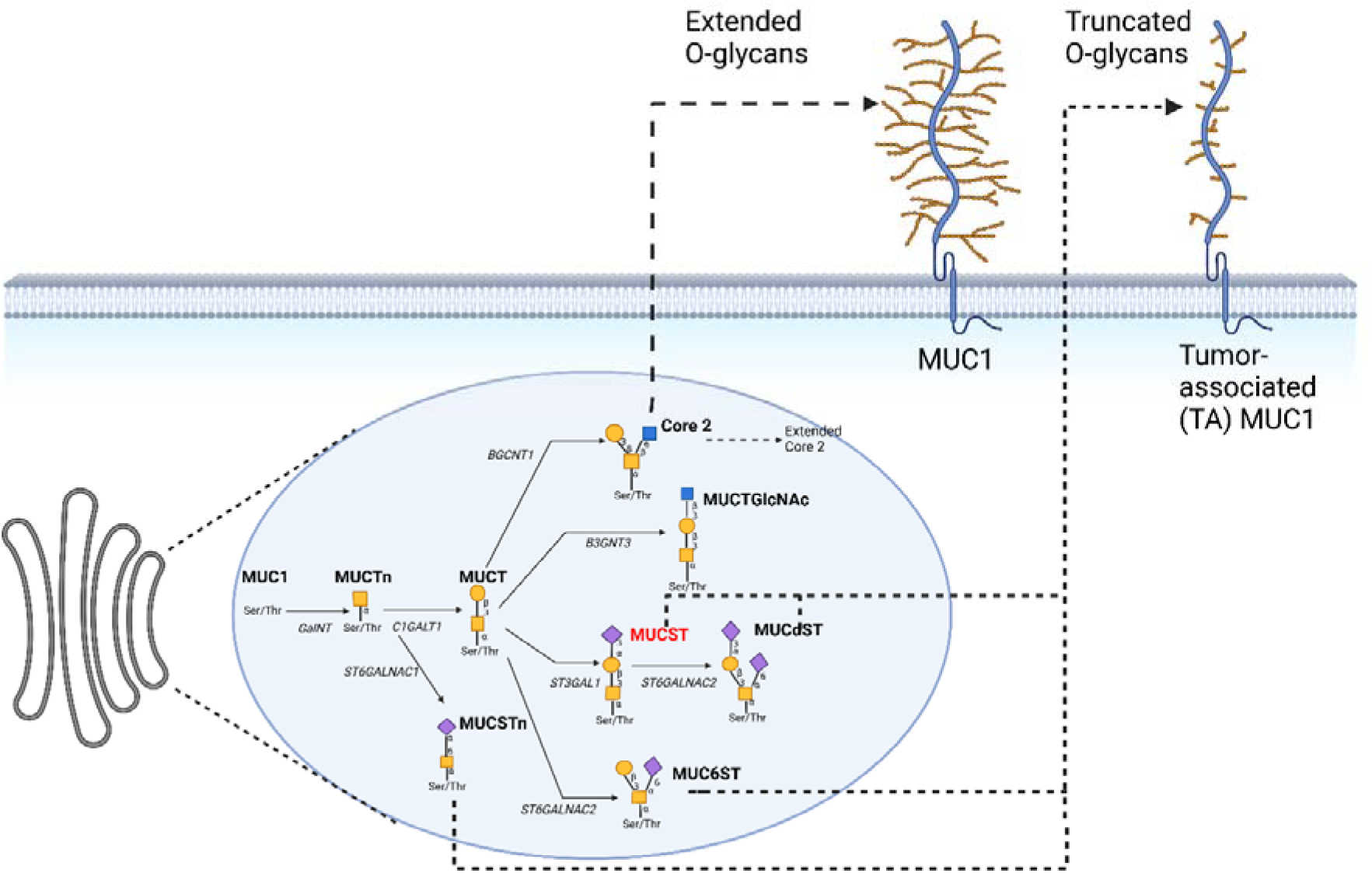
A schematic of MUC1 O-glycan synthesis in the Golgi apparatus. The extracellular domain of MUC1 bears extended core 2 glycans in a healthy state, while the tumour-associated MUC1 is characterized by truncated and sialic acid-terminated core 1 O-glycans. The O-glycan epitope symbols follow the SNFG (Symbol Nomenclature for Glycans) system (PMID 26543186, Glycobiology 25: 1323–1324, 2015) details at NCBI.

Several studies have uncovered the interactions between MUC1 O-glycan epitopes and various immune cells, such as natural killer cells (NKs) (Zhao et al. 2010), dendritic cells (DCs)(Monti et al. 2004; Napoletano et al. 2007), and macrophages. A survey of these studies reveals that the specificity of immune cell receptors towards different MUC1 O-glycan epitopes underlies the differences in immune response outcomes (Napoletano et al. 2012; Rughetti et al. 2005). A differentiation in breast cancer macrophages, identified as tumour-associated (TAMs), was uncovered following the hypersialylated core-1 epitope of MUC1 interacting with the macrophage (Beatson et al. 2016). In the study, the binding of sialyl-T antigen (Neu5Acα1-3Galβ1-3GalNAcα-O-Ser/Thr) bearing MUC1 (MUCST) by the sialic acid-binding immunoglobulin-type lectin (Siglec)-9 promoted a TAM-like phenotype, leading to increases in several cytokines, chemokines, and growth factors responsible chronic inflammation, signal activation and metastasis (Beatson et al. 2020).

Within the altered macrophage secretome, the C-X-C motif chemokine 5 (CXCL5) stands out for its contribution to poor prognosis and metastasis. The 2022 review by Deng et al. considers CXCL5 “A coachman to drive cancer progression” and provides a comprehensive list of studies highlighting the deleterious effects of CXCL5-binding by the C-X-C motif chemokine receptor 2 (CXCR2) on JNK, MAPK, PI3K/AKT, and STAT3 signalling cascades (Deng et al. 2022). In breast cancer, the CXCL5/CXCR2 axis and subsequent signal activation lead to cancer progression and epithelial-to-mesenchymal transition (EMT) (Hsu et al. 2013; Li et al. 2020).

Interfacing MUC1 O-glycosylation to its downstream biological networks presents several challenges. Most important is a lack of sufficient and accurate kinetics data.

However, biologically informed assumptions can be used to generate proof-of-concept models to provide mechanistic insight into cancer network behaviour, druggable targets and their inhibition. Given the importance of MUC1 O-glycosylation and CXCL5 overexpression in cancer research, and the therapeutic potential of sialyltransferase inhibition, the present study proposes a mathematical model connecting aberrant MUC1 O-glycan synthesis with MUCST-mediated CXCL5 production in macrophages. By linking the two networks to a third network, which represents the impact of cytokine/chemokine binding on signal activation, we provide a framework that correlates MUC1 O-glycosylation to cancer cell phenotypes. CytoCopasi aids the comparison of the dynamics of CXCL5 production and signal activation upon normal, cancerous (luminal A), and ST3Gal-I-inhibited MUC1 O-glycosylation.

## Results

### Comparative Analysis Between Normal and Luminal A- differentiated Glycosylation Networks

While healthy and luminal A-type MUC1 glycosylation networks are the same their routes to different metabolite concentrations stem from differences in their initial MUC1 concentrations and glycosyltransferase activities. The models described in the previous section were used to simulate normal and luminal A conditions in CytoCopasi. The alterations in metabolite concentration levels in luminal A-type O-glycosylation compared to normal glycosylation were obtained using CytoCopasi’s comparative simulation tools, yielding the Cytoscape network view (Fig. 2A). Upregulation and downregulation are depicted in red and cyan, respectively, while the node size corresponds to the magnitude of change. The node for non-glycosylated MUC1 was excluded in the comparative simulation.

**Fig 2:**
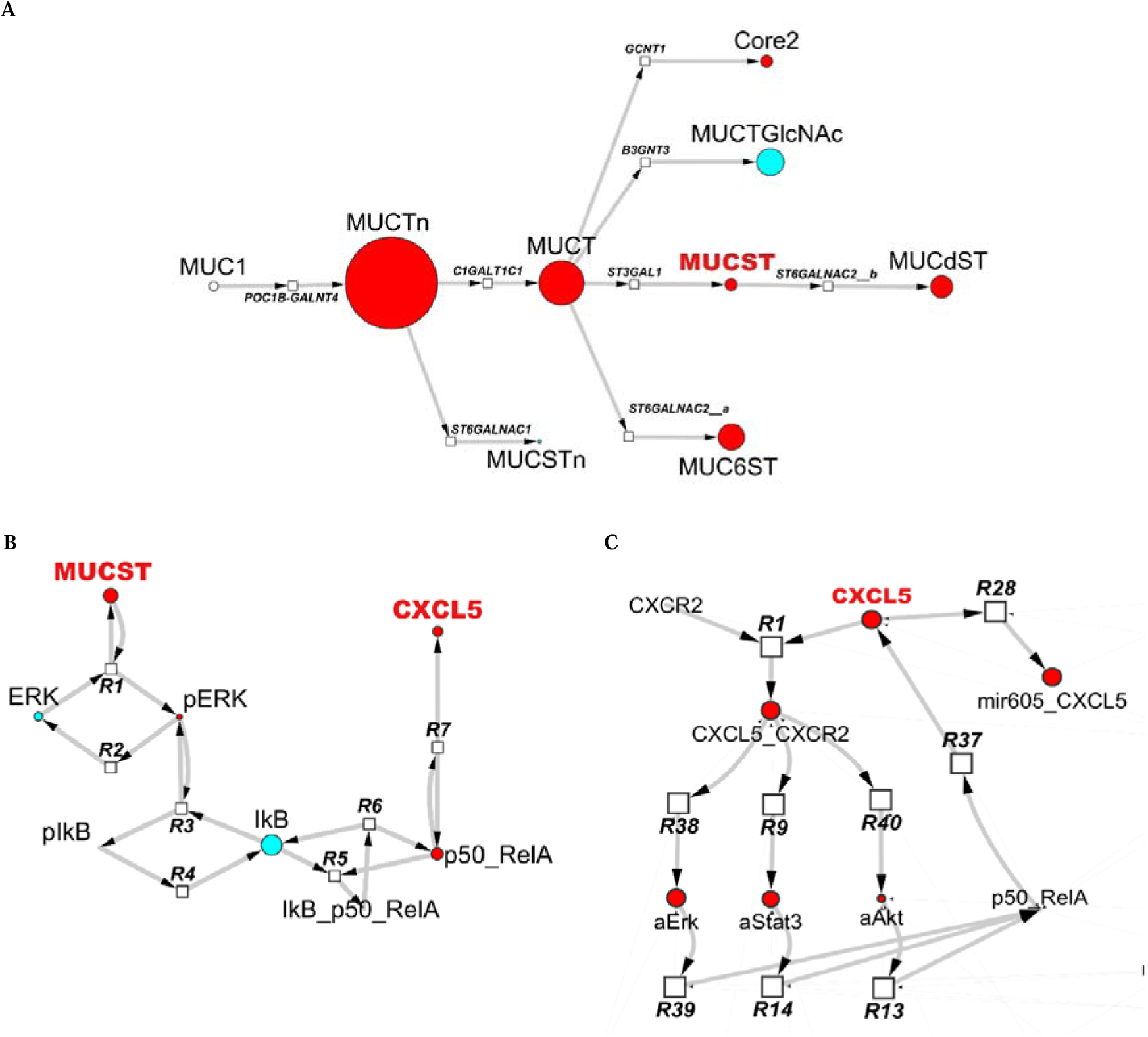
The Impact of MUC1 overexpression combined with differential luminal A glycogene expression on alterations in the MUC1 O-glycome after 80 hours. (A) An overall increase of glycan epitopes is observed, especially with the Tn- and T- antigens. After the simulation, MUCST concentration is 32% higher in the luminal A-representing network than in the healthy glycosylation network; however, the upregulation in MUC6ST (68%) and MUCdST (58%) is much higher due to increased ST6GALNAC2 activity. (B) The downstream effects of 32% MUCST upregulation in luminal A compared to normal glycosylation. The metabolite alterations and CXCL5 upregulation in the macrophage network after 80 hours. The Erk pathway is shifted towards phosphorylation, while the NF-kB activation is upregulated, leading to a 23% upregulation in CXCL5 production. (C) The downstream effects of luminal A-associated CXCL5 production on signal transduction inside the tumour cell after 80 hours. An overall increase in Stat3, Erk, and Akt pathways can be observed.

The results for core 2, and the four sialylated epitopes (Table 1a, Table S6) show a significant upregulation in MUC6St and MUCdST concentrations and smaller upregulation in MUCST concentration at 80 hours. This indicates a clear shift towards tumour-associated core 1-glycan biosynthesis in luminal A breast cancer compared to MUC1 in healthy breast epithelial tissue. The network demonstrates a 32% upregulation of MUCST, which, albeit significant, is smaller than the values for the MUC6ST and MUCdST, ranging between 55- 70%. This can be attributed to the simultaneous overexpression of ST3GAL1 and ST6GALNAC2. Although the increased ST3GAL1 activity brings about an enhanced rate in MUCST synthesis, this is counteracted by the increased enzyme activity of ST6GALNAC2 partaking in two reactions. The first reaction, denoted by *ST6GALNAC2_a* catalyses the α2,6 sialylation of the MUC1 T-antigen GalNAc residue, thereby limiting the amount of MUC1 T-antigen available for α-2,3-Sialylation at the Gal residue. The second reaction, denoted by *ST6GALNAC2_b*, catalyses the α2,6 sialylation of the GalNAc residue on MUCST, hence resulting in MUCST consumption.

**Table 1:** Perturbations in luminal A compared to healthy networks. a) Luminal A glycosylation fold changes in Core 1 epitopes and Core 2, compared to normal glycosylation at t= 288000 s. b) Impact of MUCST Upregulation in luminal A on Monocyte Metabolites at t = 288000 s. c) Impact of CXCL5 Upregulation caused in luminal A on Intracellular Signalling at t= 288000 s.

| Metabolite Name | Normal (mM) | Luminal A (mM) | Change% (Luminal A vs Normal) |
| --- | --- | --- | --- |
| MUC6ST | 0.92 | 1.55 | 68.05 ↑ |
| MUCdST | 0.51 | 0.81 | 57.71 ↑ |
| Core2 | 2.20 | 2.91 | 32.36 ↑ |
| MUCST | 1.02 | 1.35 | 32.12 ↑ |
| MUCSTn | 0.00103 | 9.20 x 10<sup>-4</sup> | 10.95 ↓ |

b)
| Metabolite | Before Perturbation (mM) | After Perturbation (mM) | Change% |
| --- | --- | --- | --- |
| IκB | 4.93 x 10<sup>-6</sup> | 2.75 x 10<sup>-6</sup> | 44.20 ↓ |
| p50_RelA | 2.31 x 10<sup>-10</sup> | 2.92 x 10<sup>-10</sup> | 26.72 ↑ |
| CXCL5 | 8.12 x 10<sup>-9</sup> | 9.98 x 10<sup>-9</sup> | 22.84 ↑ |
| ERK | 4.50 x 10<sup>-5</sup> | 3.60 x 10<sup>-5</sup> | 20.02 ↓ |
| pERK | 6.60 x 10<sup>-5</sup> | 7.50 x 10<sup>-5</sup> | 13.66 ↑ |

c)
| Metabolite | Before Perturbation (mM) | After Perturbation (mM) | Change% |
| --- | --- | --- | --- |
| aErk | $5.32 \times 10^{-5}$ | $6.6 \times 10^{-5}$ | 22.52 ↑ |
| mir605_CXCL5 | 0.00323 | 0.00396 | 22.38 ↑ |
| CXCL5_CXCR2 | 0.00323 | 0.00396 | 22.38 ↑ |
| aStat3 | $5.85 \times 10^{-5}$ | $7.1 \times 10^{-5}$ | 20.50 ↑ |
| aAkt | $1.12 \times 10^{-4}$ | $1.24 \times 10^{-4}$ | 10.80 ↑ |

**Table 2:**
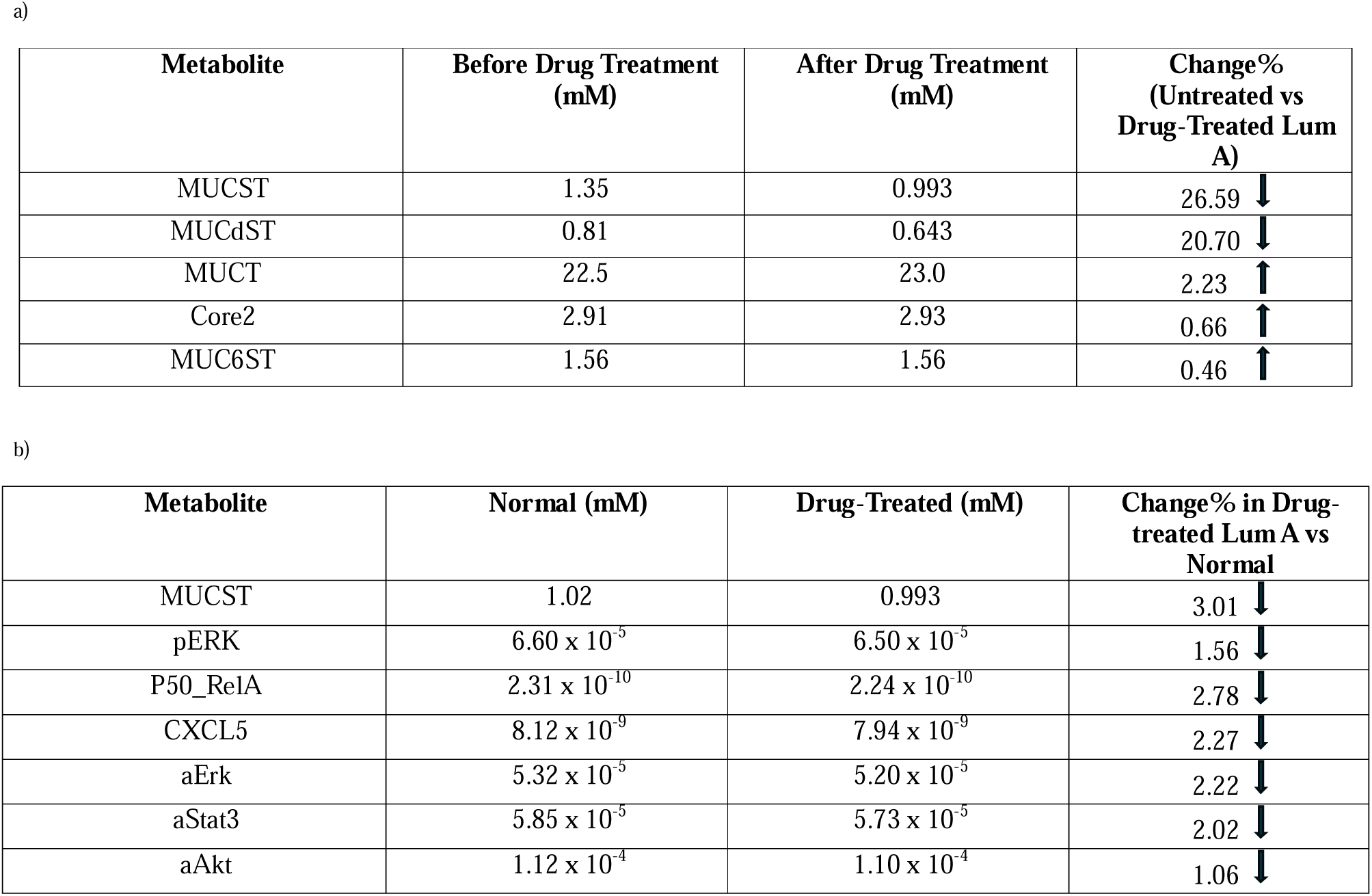
Changes in Network Metabolite Levels Upon ST3Gal-I inhibition with Soyasaponin-I. a) Direct effects of increasing initial Soyasaponin I concentration from 0 to10 µM on luminal A at t= 288000 s. b) Fold changes in key metabolite levels for drug-treated luminal A compared to normal metabolite levels at t= 288000 s.

Nevertheless, a 32% upregulation of MUCST in luminal A breast cancer is substantial enough to investigate the influence of increased MUCST binding on the CXCL5 synthesis pathway in monocyte-derived macrophages.

Metabolomic alterations in the macrophage were achieved by altering the initial concentration of MUCST. CytoCopasi’s sequential perturbation algorithm can directly extract the fold increase of MUCST from aberrant glycosylation and use this value as a factor to set the initial and perturbed values of MUCST concentration at the beginning of the macrophage network simulation. Subsequently, two versions of the macrophage network are simulated for 80 hours, one with the original value for initial MUCST concentration *(0.111 µM)* and one with a perturbed initial concentration *(0.111 x (1+0.32) = 0.147 µM)*.

As illustrated in Fig.2B, and summarized in Table 1b and Table S7, a 32% increase of the initial MUCST concentration occasions upregulations in pERK and p50_RelA (Active NF-kB) by 13.7 and 26.7%, respectively. CXCL5 is upregulated by 23% compared to its original value in the case of normal-like O-glycosylation. This increase can be attributed to a shift of the ERK pathway towards phosphorylation, causing a substantial decrease in active IkB and leaving room for further p50_RelA nuclear translocation.

The perturbation of the intercellular signalling network was induced by the upregulation of CXCL5 obtained from the macrophage network. The model was simulated for 3.3 days to map out the immediate effects of excessive CXCL5 in the TME. A substantial overactivation of the Erk (22.3%) and Stat3 (20.3%) pathways (Fig. 2C, Tables 1c and S8), and to a smaller extent, the Akt pathway (10.7%) was observed. The smaller upregulation of Akt can be explained by the number of receptor-ligand complexes influencing each pathway in the network. Erk is added to the original model and is only influenced by the CXCL5- CXCR2 complex which is consistent with reports on the significance of CXCL5 in breast cancer mainly points to the MAPK signalling cascade (Hsu et al. 2013; Zhao et al. 2017). In contrast, the Akt amount is determined not only by the CXCR2/CXCL5 axis but also by IL8/CXCR1 and HER2/EGFR axes. Thus, the increase in CXCR2/CXCL5 association would be expected to have a less profound effect on the net upregulation of Akt than it does on Stat3 and Erk.

A 22% increase in CXCL5 and CXCL5-CXCR2 after 3.3 days clearly indicates the accumulation of the chemokine in both free and complex forms. The accumulation comes about despite the increase in CXCL5 inhibition by miR605 seen from the upregulation of miR605_CXCL5 (Fig. 2C). The excessive chemokine in the TME may therefore overwhelm the innate mechanisms of chemokine inhibition in the tumour cell. This implies a possible target for anti-cancer therapeutic inhibition.

### Investigation of the effects of 10 µM Soyasaponin I on Luminal A-differentiated Glycosylation and Downstream Networks

The inhibitory effects of Soyasaponin I were evaluated based on the extent to which the drug could reduce MUCST upregulation at t= 80 hours compared to normal glycosylation.

The effects of 10 µM Soyasaponin introduction on the cancerous network (Table 3a, Tables S9-S11) results in a 26.6% decrease in MUCST levels. The largest change is observed for MUCST in line with the expectations since the drug primarily targets ST3Gal1, the enzyme catalysing MUCST production. The substantial decrease in MUCST leads to an associated decrease in MUCdST by 20.7%. In addition, there is an increase of 2.2% in MUCT levels at t=80 hours is observed, despite the inhibitor tangentially acting on C1GALT1 due to a weak binding affinity of the inhibitor to the enzyme. Furthermore, increases below 2% do occur with other core structures, such as core 2, MUC6ST, and MUCTGlcNAc.

To assess the extent to which inhibition by Soyasaponin restores the three networks to their original states, the drug-treated cancerous glycosylation network was compared to the normal glycosylation network. From this, Soyasaponin appears to be reasonably effective if the MUCST percentage changes between the drug-treated and the normal networks are effective downstream. Indeed, Soyasaponin I achieves a decline in CXCL5 synthesis in the macrophage network by 2.3% (Table 3b. Tables S12-S14, and Fig. 3A). Meanwhile, the overall reduction in signal activation for Erk, Stat3, and Akt ranges between 1.1-2.2%.

**Fig. 3:**
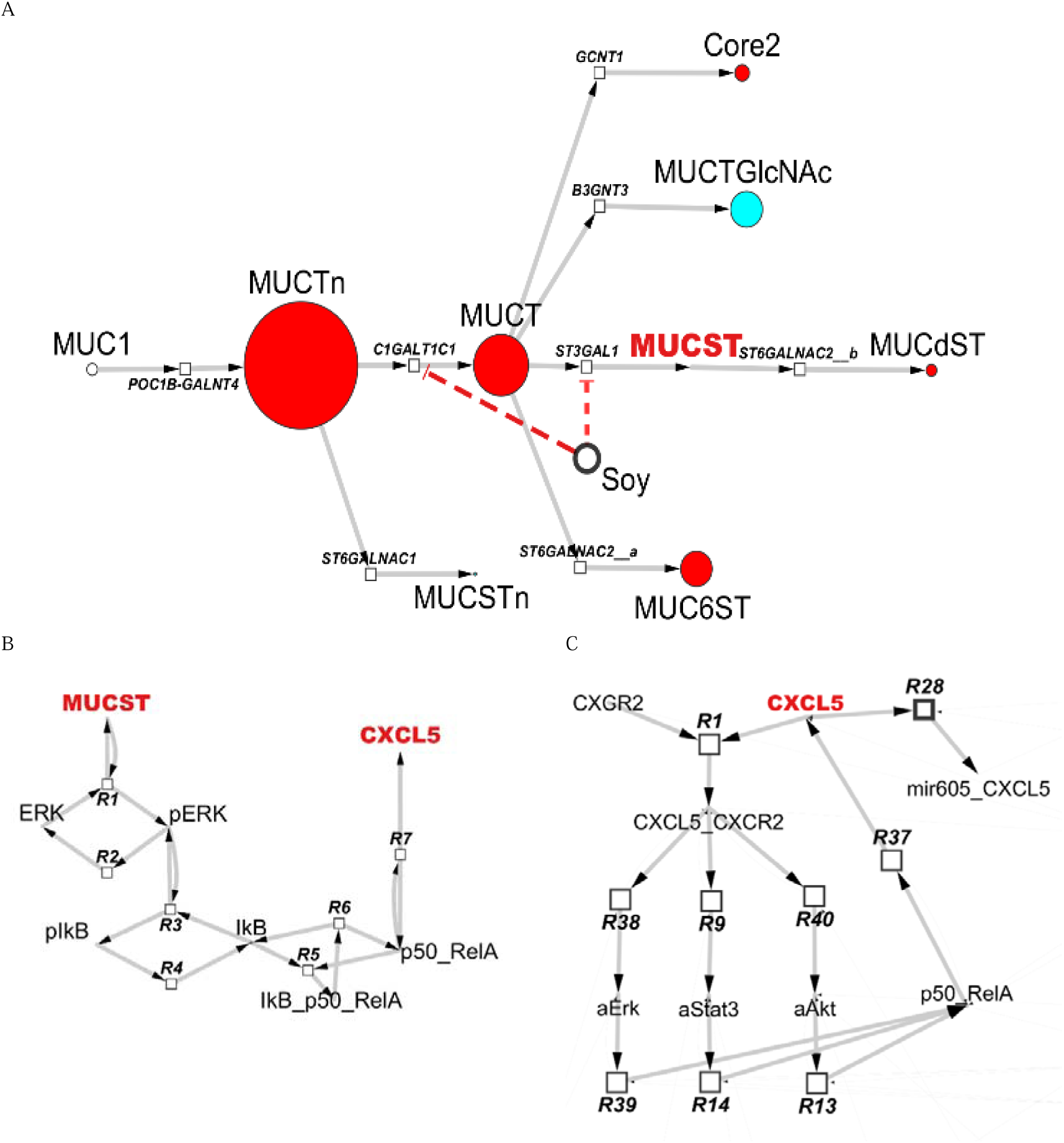
Evaluation of the ST3Gal1 inhibitor Soyasaponin I on luminal A inferred from comparison to the normal state. (A) Compared to the normal glycosylation network, MUCST transient concentration at 80 hours is 3% less in the 10 µM Soyasaponin-treated luminal A cancer network. On the other hand, C1GalT1 inhibition does not suffice to restore the MUCT concentration to its normal value. (B) 10 µM Soyasaponin is sufficient to maintain ERK and p50_RelA activity near the original levels. (C) Thus, 10 µM Soyasaponin suffices to abrogate signal activation in the luminal A breast cancer cell.

However, the Cytoscape nodes for the macrophage and intercellular signalling network are not easily seen (Fig. 3B-C) as the magnitude of metabolite percentage change to the corresponding node is mapped. Overall, the comparative simulation between drug-treated luminal A and normal glycosylation shows that 10 µM Soyasaponin may be sufficient to completely reverse aberrant CXCL5 production in the macrophage and signal activation in the breast cancer cell.

## Discussion

We advance systems glycobiology beyond models of biosynthesis that optimize for a specific recombinant glycoprotein. Our present model lays the groundwork for biochemical reaction networks to simulate the effect of changes in glycosylation on essential cellular processes. Here, we link aberrant glycosylation to the TAM secretome and enhanced signal transduction, which are biochemical hallmarks of breast cancer. The visual analytics through CytoCopasi workflows provide a comparative study of normal and cancerous glycosylation. The distinct O-glycomes of normal epithelial cells and luminal A-type cells show their differing interactions with macrophages. In the latter, the resultant MUCST directly binds to siglec 9 on the macrophage, leading to the release of the pro-inflammatory CXCL5, which overactivates signal transduction in the tumour cell. This pilot study proposes an ODE-based model delineating the downstream effects of glycosylation in cancer progression.

An established limitation of chemical systems glycobiology models is the lack of organism-consistent glycoenzyme kinetics, which prevents accurate parameterization.

However, quantitative omics data, such as differential gene expression from the TCGA, can be used for perturbations and model fitting that result in proof-of-concept models. A drawback of TCGA-based perturbation is the assumption that altered glycosyltransferase gene expression is completely translated into the synthesis of MUC1 O-glycans, discounting the impact of these alterations on other glycoproteins. Nevertheless, this data can provide the causal link between glycogene overexpression and MUC1-related tumourigenicity. For example, Picco et al. showed that mice overexpressing ST3Gal1 in the presence of the MUC1 promoter displayed earlier tumour development than the wild-type control, producing higher levels of tumour-promoting cytokines (Picco et al. 2010). Interestingly, ST3Gal1 overexpression did not enhance the interaction of the MUC1 cytoplasmic tail with the proto-oncogene tyrosine-protein kinase (c-Src), even though this interaction is a strong indicator of tumour development in the mammary gland (Al Masri and Gendler 2005). Thus, the role of ST3Gal1 overexpression in mammary tumour development is likely related to MUCST abundance in the extracellular domain engaged by the immune cells in the TME.

Nevertheless, TCGA data must be used cautiously while parameterizing glycosylation networks, as the evidence of glycogene overexpression and glycan epitope abundance may not always agree or be available. For instance, ST6GalNAc1 was underexpressed in luminal A according to TCGA, leading to downregulated MUCSTn in the CytoCopasi network.

Indeed, ST6GalNAc1 under expression can be positively correlated with better survival rates in breast cancer (Patani et al. 2008). In another instance, higher levels of STn have been associated with increased PD-L1 expression and poor prognosis in breast cancer patients (Xu et al. 2021), while ST6GalNAc-I overexpression in MCF-7 (luminal A) and BT474 (luminal B) increased the invasive capabilities on cancer cells (Luo et al. 2023). Collectively taken, these studies point to the dichotomy between glycosyltransferase expression data and the role of glycosyltransferases in driving cancerous phenotypes. Understanding the MUC1 epitope distribution in this context requires standardized metabolomics and enzyme kinetics studies. To address the dearth of consistent data, we have developed a universal glycosyltransferase continuous (UGC) assay for accurate and organism-consistent glycosyltransferase parameter measurement (Nashed and Naidoo 2024). In an illustrative study, kinetic parameters for three glycosyltransferases C1GalT1, ST3Gal1, and Fucosyltransferase I (Fut1) were estimated.

These enzyme functional parameters K_m_ and V_max_ were not used here as the rest of the enzymes’ kinetics data are not yet available from this platform.

A key objective in ODE-based models is to design frameworks that evaluate the degree to which therapeutics can return diseased systems to normal organism function. Here we considered the effect of the sialyltransferase (ST3Gal1) inhibitor Soyasaponin-I on the luminal A breast cancer subclass. Soyasaponin-derived inhibitors were discovered to suppress the FAK/paxillin pathway and circumvent angiogenesis and metastasis of cancer cell lines (Chen et al. 2011). The interaction of FAK-Paxillin with Erk and Stat3 signal transduction networks has central implications for cell growth, proliferation, and wound healing in healthy cells (Kafi et al. 2020; Liu et al. 2002; Teranishi et al. 2009), and aggressive cancer phenotypes (Pribic and Brazill 2012; Silver et al. 2004; Sp et al. 2017; Wen et al. 2020).

Therefore, the FAK/Paxilin axis might be incorporated into future models to link ST3Gal1 expression to a more elaborate signalling network. Furthermore, the upregulations in MUCT and the MUC6ST caused by ST3Gal1 inhibition highlight the importance of dosing, as excessive Soyasaponin concentration might shift the system towards the tumour-associated MUC6ST (Dziadek et al. 2006).

Taken together, the ST3Gal1 inhibition proposed in the present model imparts possible mechanisms of drug action through immunomodulation, championing drug discovery studies for sialyltransferase inhibition and addressing the need for a combinatorial inhibition to prevent tumour-associated O-glycan biosynthesis in its entirety.

While the present model assumed that the initial MUCST upregulation was entirely reflected in the macrophage network, future models need to consider regulatory factors, such as MUCST diffusion through the ECM and Siglec-9 ··· MUCST binding kinetics, to accurately quantify the increase in the MUCST input to the macrophage network. Furthermore, while the initial concentrations of the non-zero metabolites were set uniformly, knowledge of their relative abundances enabled by MS-based proteomics could yield a more realistic representation of the starting conditions. ELISA and q-PCR experiments probing ERK and NF-KB activation in response to MUCST binding by Siglec 9 can improve parameter estimation. Finally, monitoring CXCL5 production in response to different MUCST concentrations could help validate the input-output correlation of the macrophage model.

## Materials and Method

### Construction of the Networks in the Tumour Macrophage Response Model

The proposed model shown in Fig. 4 consists of three networks. It is initiated with the O-glycosylation of MUC1 in the Golgi of a breast epithelial cell, resulting in core 1 epitopes, including MUCST, and core 2. MUCST shed into the TME is recognized by Siglec9 on the macrophage surface, which promotes CXCL5 production through MEK/ERK and NF-kB. CXCL5 gets secreted into the TME and is bound by CXCR2 on the cell surface, inducing signal activation, and autocrine CXCL5 production.

**Fig. 4:**
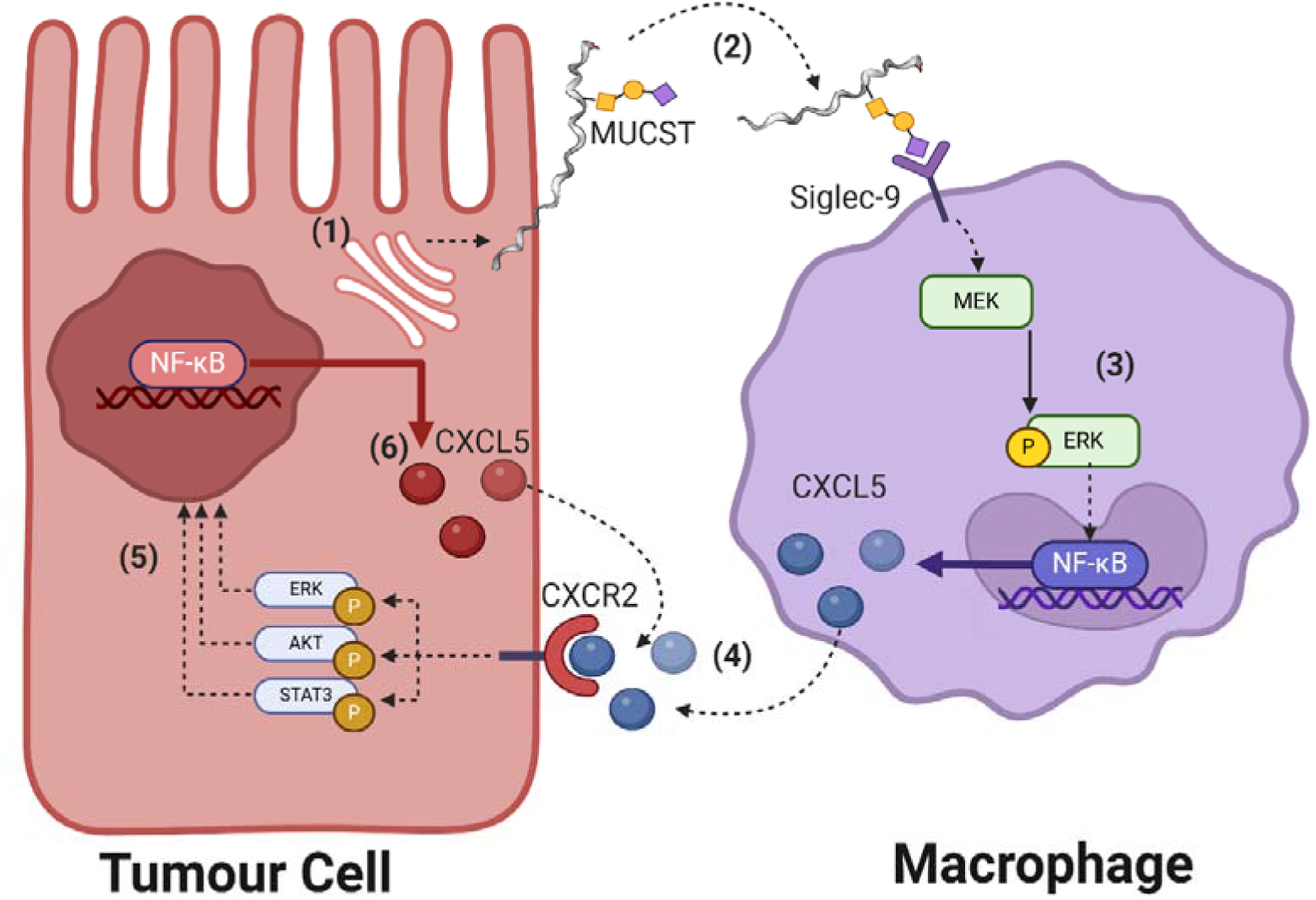
The sequence of the interplay between a tumour cell and a macrophage. (1) Aberrant glycosylation in the Golgi of the tumour cell shifts the O-glycome of MUC1 towards core 1, upregulating MUCST amongst other sialylated glycan epitopes. (2) MUCST shed from the tumour cell surface is bound by Siglec-9 on the macrophage (3) Bound siglec-9 activates ERK activation by MEK, and phosphorylated (active) Erk facilitates CXCL5 production through NF-kB activation. (4) CXCL5 secreted into the TME is bound by the CXCR2 on the tumour cell surface. (5) The CXCL5/CXCR2 axis activates Erk, Akt, and Stat3, giving rise to autocrine CXCL5 synthesis, (6) Autocrine CXCL5 binds CXCR2 and creates a positive feedback loop that sustains signal activation and CXCL5 amounts.

### MUC1 O-Glycosylation in the Breast Cancer Cell

The network structure is based on the Mucin type O-Glycan Biosynthesis network on KEGG (ID: hsa00512), which can be directly imported into CytoCopasi as an ODE-based SBML file. Core 3 and 4 structures were removed to maintain the focus on the two abundant cores 1 and 2. Assuming an excess amount of sugar donor, each of the reactions was re- written in the following form to adapt to simple Michaelis-Menten kinetics:

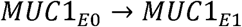

Where *MUC*1*_E_*_0_ is the monosaccharide acceptor and MUC1 target epitope, while *MUC*1*_E_*_1_ is the new epitope following the glycosidic bond formation with a monosaccharide gained from a sugar-phosphate donor through glycosyltransferase catalysis.

The resulting network, shown in Fig. 5A, starts with the synthesis of the Tn-antigen MUCTn from the MUC1 peptide by polypeptidyl GalNAc transferase (GALNT1). Core 1 β1,3 – galactosyltransferase (C1GalT1) catalyses the addition of a galactose residue to the MUCTn to form T antigen (MUCT). Further addition of GlcNAc by core 2 β1 → 6 N-acetylglucosaminyltransferases (C2GnTs) leads to the core 2 starting structure. The model also considers the linear GlcNAc extension of Core 1, catalysed by β1,3-N-acetylglucosaminyltransferase-3 (B3GNT3). Tumour-associated glycan epitopes contain sialic acid (Neu5Ac) as their terminal residues. α2,6 sialyltransferase (ST6GalNAc1) and β- Galactoside α-2,3-Sialyltransferase 1 (ST3Gal1) are responsible for the formation of Sialyl-Tn antigen (MUCSTn) and Sialyl-T antigen (MUCST), respectively. ST6GalNAc2 adds Neu5Ac to the GalNAc residue of the T- and ST-antigens to form MUC6ST and MUCdST.

**Fig. 5:**
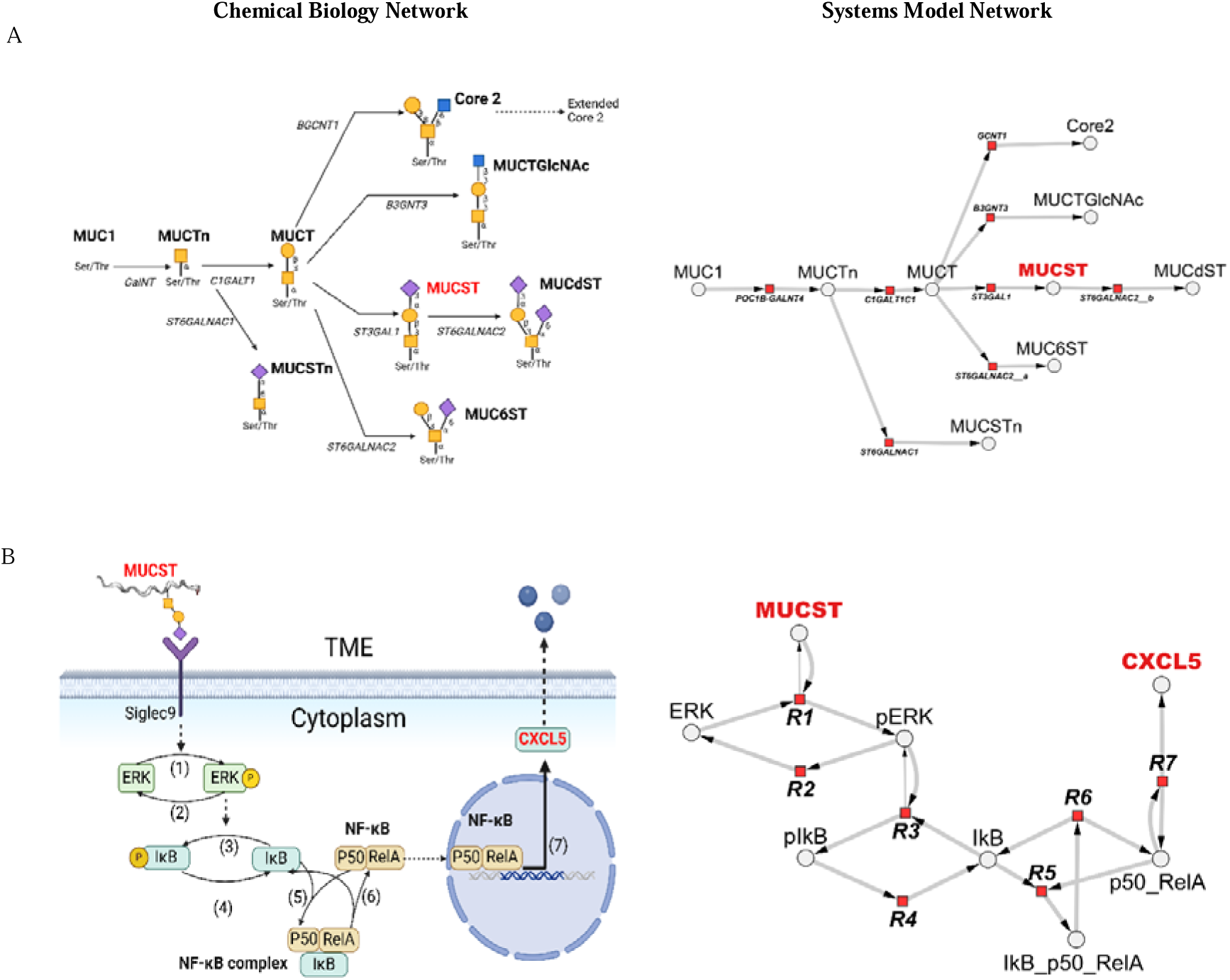
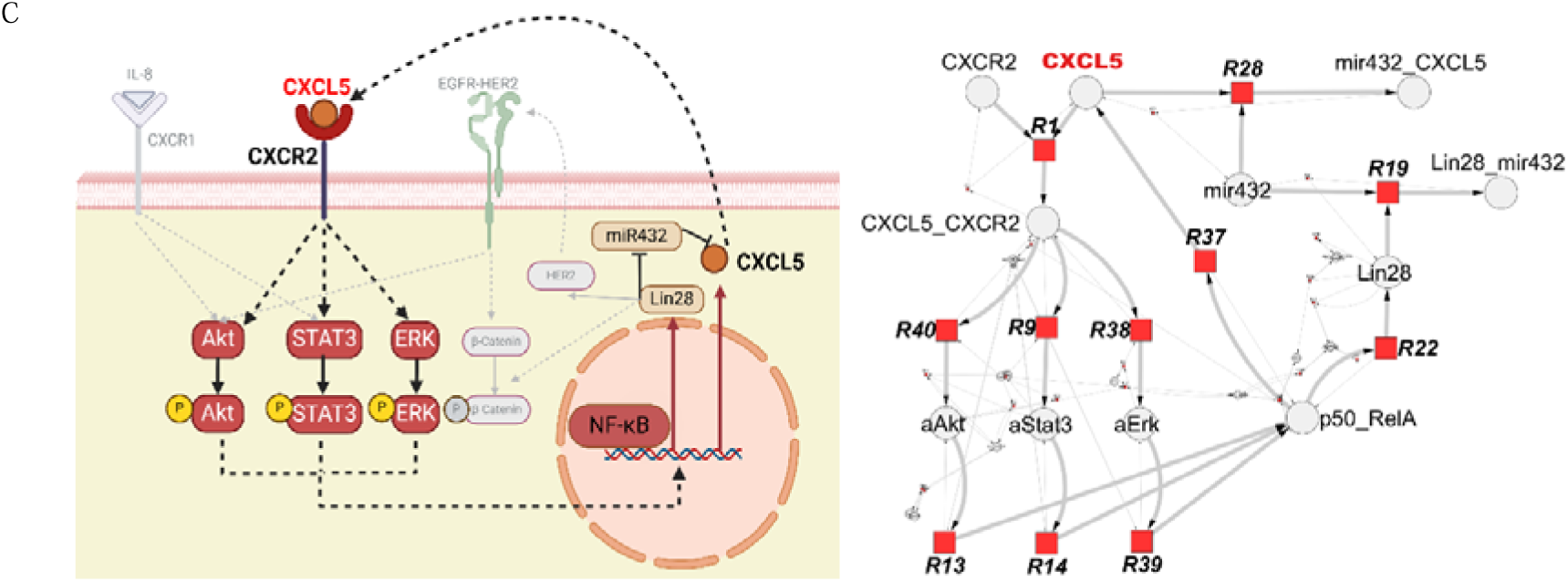
Schematic of a chemical biological network (left) and CytoCopasi systems biology network (right) views of the constructed networks. (A) MUC1-type O-glycosylation in breast tissue contains MUC1, MUC1-Tn, and MUC1-T antigens as well as the core 2 epitope and core 1 structures MUC1- STn, MUC1-ST (MUCST), MUC1-6ST, MUC1-dST, and extended Core 1. The O-glycan epitope symbols follow the SNFG (Symbol Nomenclature for Glycans) system (PMID 26543186, Glycobiology 25: 1323–1324, 2015) details at NCBI (B) The reaction network depicting the set of events from MUCST-Siglec-9 binding through Erk stimulation, IkB phosphorylation, NF-kB activation (C) The positive feedback loop of CXCL5 production. CXCL5-CXCR2 interaction promotes the activation of Akt, Stat3, and Erk, all of which contribute to the activation of NF-kB. As a result, more CXCL5 is synthesized within the cell, accumulating in the TME for further CXCR2 binding.

The initial concentration of the MUC1 peptide was set to remain constant at 1.88 mM based on reported values for milk mucin (Peterson et al. 1998). The other metabolites, i.e., the glycan epitopes were set to 0.

The K_m_ for the enzymes were extracted from the BRaunschweig ENzyme DAtabase (BRENDA) (Schomburg et al. 2002) using CytoCopasi (Kaya and Naidoo 2023). The acceptor-specific K_m_ value was used whenever possible. Due to the absence of data for ST6GalNAc2, the K_m_ value for ST6GalNAc1 was set for the three ST6GalNAc-driven reactions.

The network was fitted to the O-glycan metabolomics of mature human milk, where the ratio of sialylated to non-sialylated mucin O-glycans was given as ∼0.24 (Lu et al. 2019). The Genetic Algorithm optimization method of Complex Pathway Simulator (COPASI) was used, with time course as the subtask (Hoops et al. 2006). The complete list of V_max_ and K_m_ values, as well as the primary references for the K_m_ values are provided in the Supplementary Document (Table S1).

### CXCL5 Production in the Monocyte-derived Macrophage

The study by Beatson et al. proposed the involvement of MEK-ERK activation in the increase in the MUCST-mediated secretion of cytokine and chemokines, including CXCL5 (Beatson et al. 2016). The underlying mechanism in macrophage secretome alterations can be inferred from known biochemical networks. Firstly, the transcription factor nuclear factor kappa B (NF-kB) plays an instrumental role in immune regulation through the synthesis of cytokines and chemokines (Zhang et al. 2021). Its heterotetrametric form, consisting of a ∼50 kD subunit (p50) and two ∼65 kD subunits (p65/RelA), NF-kB can rapidly bind its cognate DNA to initiate transcription (Kawakami et al. 1988). NF-kB activity is inhibited by the binding of the kappa B protein (IkB), which prevents its translocation into the nucleus (Baeuerle and Baltimore 1988). The interplay between NF-kB and IkB is modulated by IKappaB kinase (IKK), which phosphorylates IkB and blocks its binding to NF-kB (Zandi et al. 1997). Furthermore, studies of inflammation and cancer implicate ERK activation as a driver of IKK activation (Suh et al. 2002; Zhao and Lee 1999). Thus, by activating IKK, ERK promotes the inhibition of IkB, which contributes to NF-kB activation. These interactions inform the construction of a network describing altered macrophage secretomes.

The monocyte model structure was constructed based on the reaction network proposed by Figueiredo et al. (Figueiredo et al. 2009), illustrating the cytokine production of macrophages upon external stimulus by *A. viteaei.* Here in this model, the external stimulus is replaced with MUCST which promotes ERK phosphorylation that in turn induces IkB phosphorylation. When non-phosphorylated, IkB forms a complex with p50_RelA, keeping it sequestered from the nucleus. IkB phosphorylation prevents complex formation, allowing the nuclear translocation of p50_RelA to drive CXCL5 synthesis. The model includes the constitutive dephosphorylation of ERK and IkB which is illustrated in a reaction network that consists of 8 metabolites and 7 reactions (Fig. 5B).

The Initial concentration of MUCST was set to 0.111 µM based on the experimental procedures provided by Beatson et al. (Beatson et al. 2020). The non-zero metabolites, ERK, IkB_p50_RelA, and IkB were uniformly set to 0.111 µM. Phosphorylated forms of ERK, IkB, and the free form of NF-kB (p50_RelA) were set to zero. Spatial evolution of the species, IkB synthesis and phosphorylated IkB degradation were excluded. Finally, the negative feedback loop between CXCL5 and ERK was excluded to replicate a sustained cancerous ERK phosphorylation (Orton et al. 2009).

Mass-action kinetics formed the foundation of the model reactions. Parameter estimation was performed using the Particle Swarm algorithm (Kennedy and Eberhart 1995).

Seven rate constants were estimated to fit the model to CXCL5 production in monocytes co-cultured with MUCST-bearing T-47D cell lines (Beatson et al. 2020). The estimation was constrained to keep the K_d_ value for the IkB_p50_RelA complex *(R6.k1/R5.k1)* within a reasonable range based on reported values (Urban and Baeuerle 1990). The complete list of reactions, rate laws, and estimated values used can be found in the Supplementary Document (Table S2).

Parameter estimation yielded an objective function of ∼10^-22^, and the fitted curve illustrated in Figure S1 coincides with the experimental data points, capturing the exponential increase in CXCL5 concentration over time. The time-course concentration profile of the model shown in Figure S1 displays a decrease in ERK and IkB, as their phosphorylated versions increase.

### Signal Activation in the Breast Cancer Cell

The final component of our model (Fig. 5C), is a modified version of the reaction network by Sehl et al. (Sehl et al. 2015), which simulates the positive feedback loop between IL-6, signal transduction, and NF-κB as the driving factor of EMT in breast cancer stem cells. This study was selected as the starting point, as it recapitulated the effects of cytokines and chemokines on intracellular signalling and transcription.

The initial model was modified to introduce CXCL5. Firstly, IL-6 was replaced with CXCL5. Instead of gp130 from the IL-6 receptor family, CXCL5 binds CXCR2. ERK and activated ERK were added following studies reporting ERK activation by CXCL5-CXCR2 binding (Hsu et al. 2013). CXCR2/CXCL5 activates STAT3, AKT, and ERK, which subsequently activate NF-κB by promoting its dissociation from the IκB. Thus, the cancer cell produces CXCL5, which becomes a ligand for CXCR2. The continuous activation of the CXCL5/CXCR2 axis causes sustained activation of the signalling pathways. CXCL5 accumulation is inhibited by miR432 instead of Let7 (Luo et al. 2021).

The cell population dynamics of epithelial and mesenchymal cells were removed to keep the analysis on a molecular level. Whereas profound changes in cell populations required simulation over years, the removal of these cells enables the observation of the immediate effects of chemokine overexpression on signal transduction and transcription. The complete list of the modified and added reactions can be found in the Supplementary Document (Table S3-S4).

### CytoCopasi Algorithm for Sequential Network Perturbations

Simulations were performed with CytoCopasi to compare signal activation in health and different cancer networks. A detailed description of the comparative simulation workflows can be found in the work reporting CytoCopasi development (Kaya and Naidoo 2023). The default CytoCopasi’s ODE solver algorithm, LSODA (Petzold 1983), was used to run simulations for 288000 seconds across all networks. The simulation length was chosen for consistency with the experimental data on CXCL5 that ran for 80 hours (Beatson et al. 2020).

The simulation of the three networks requires multiple perturbations, starting with aberrant MUC1 glycosylation. Upregulated MUCST as observed in cancer, is used as input for the monocyte/macrophage network. Following this, the CXCL5 output of the second network becomes the input for the intracellular signalling network. Consequently, three models are sequentially perturbed to model the impact of the aberrant glycosylation on intracellular signalling. A CytoCopasi module was developed to streamline the interlocking of these perturbations, which automatically transfers in the metabolite fold changes from a precursor simulation and to perturb the initial values of the corresponding metabolites in the subsequent network, with the proviso that at least one metabolite is common between the interlocking networks. Once percentage concentration changes (PC) are generated from the first network simulation, the user chooses the follow-on network, selects the bridging metabolite(s) and specifies the subtask as time-course and its duration. For each selected metabolite *A*, the algorithm extracts the PC value from the current network and defines the unperturbed and perturbed values in the next network as follows:

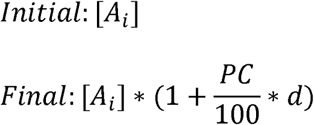

Where *d* is either 1 or -1 depending on whether *A_i_* was upregulated or downregulated in the initial network.

This algorithm is implemented twice in a workflow (Fig. 6A) that is initiated with a comparative simulation of the MUC1 O-glycosylation between normal and luminal A states for 80 hours (288000 seconds), where the luminal A model was constructed using differential gene expression data from The Cancer Genome Atlas (TCGA) (Tomczak et al. 2015). To achieve this the initial MUC1 concentration was multiplied by the fold change of MUC1 expression and V_max_ values of glycosyltransferases were multiplied by the fold changes in the corresponding glycogene expression. The TCGA data and its incorporation in the luminal A glycosylation network are provided in Table S5. We know that the relationship between differential gene expression and enzymatic expression may not be directly correlated (Pothukuchi et al. 2019). However, it is known that sialyltransferase overexpression in the presence of the MUC1 promoter accelerates tumour growth in mammary glands, and so validates the use of TCGA data to inform the construction of tumour-associated glycosylation networks (Picco et al. 2010).

**Fig. 6:**
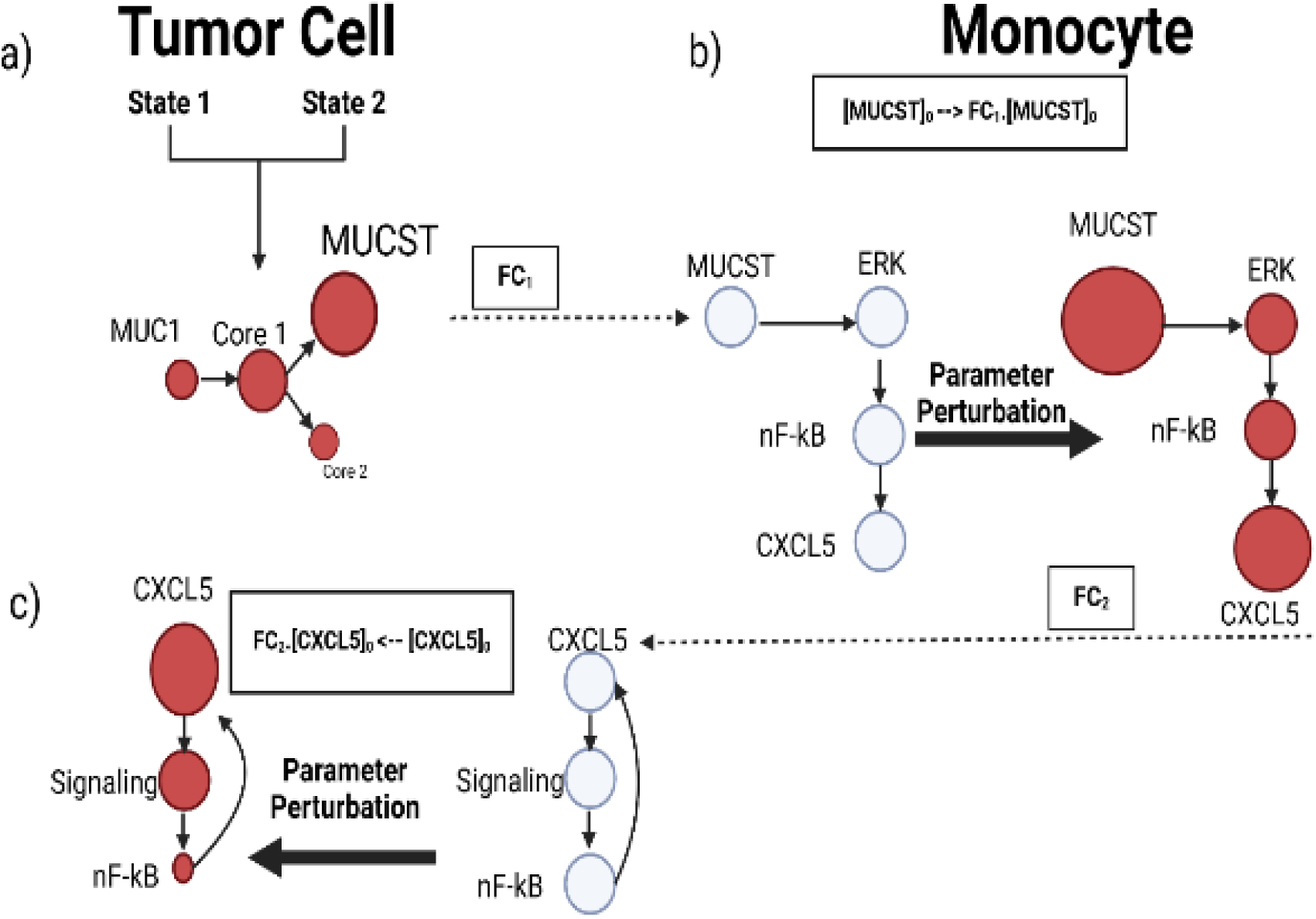
A simplified depiction of the algorithm used to perturb models sequentially. (A) A comparative simulation of two Mucin O-glycosylation networks reveals an upregulation of MUCST. (B) The fold change of MUCST is used to perturb the initial concentration of MUCST in the monocyte network, driving an increase in ERK and NF-kB activation, and the eventual CXCL5 production. (C) The fold change of CXCL5 is used to perturb the initial value of CXCL5 in the intracellular signalling network, leading to an overactivation of the signalling cascades.

Fold change values were generated for glycan epitopes between luminal A and normal glycosylation. The initial concentration of the MUCST in the macrophage network is then perturbed by the output altered value from the glycosylation network (Fig. 6B). An 80-hour simulation of this network leads to alterations in metabolite transient concentrations. The second perturbation is run by selecting an intracellular signalling network, where the fold change in monocyte-CXCL5 is reflected in the initial CXCL5 concentration (Fig. 6C).

### Inhibition Kinetics Studies

Soyasaponin-I was introduced as an ST3Gal1 inhibitor with an IC_50_ value of 0.031 mM for ST3Gal1 and 0.368 mM for C1GalT1 (Nashed and Naidoo 2024). A drug-treated version of the luminal A-type glycosylation network was created, containing the same cancerous perturbations and, additionally, 0.010 mM Soyasaponin I as a competitive inhibitor for *ST3GAL1* and *C1GALT1* (Wu et al. 2001). A new rate law function in terms of IC_50_ for competitive inhibition was created in CytoCopasi by solving for *K_i_* in the equation (Wei et al. 2007)

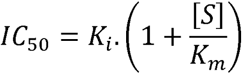

And substituting for *K_i_* in the competitive inhibition rate law to yield.

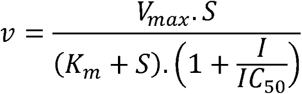

Where *S* and *I* refer to the transient substrate and inhibitor concentrations, respectively.

The comparative analysis of ST3Gal1 inhibition involved two comparisons: Comparing the drug-treated cancer glycosylation network to its non-treated version would allow the monitoring of the direct effects of the drug on MUCST downregulation and the downstream networks. On the other hand, comparing the drug-treated cancerous glycosylation to the original healthy network reveals whether the drug is sufficient to restore normal MUCST levels. We use as a benchmark the performance of an ideal inhibitor that minimizes MUCST upregulation resulting in unperturbed macrophage and intracellular signalling networks.

## Supporting information

Details of model construction and Tables listing reactions parameters and formulas

## Acknowledgements

The authors thank Dr Richard Beatson, Kings College London, for providing insight into the experimental data used in model fitting. The authors also thank Dr. Pedro Mendes, UConn School of Medicine, and Dr. Alfonso Landeros, UC Riverside for their suggestions during model construction. EK Thanks the Scientific Computing Research Unit (SCRU) for graduate fellowship funding. We thank the Centre for High-Performance Computing (CHEM0840) for providing additional computing resources.

## Abbreviations

MUCTn: GalNAcα–O-Ser/Thr
MUCT: Galβ1-3GalNAcα-O-Ser/Thr
MUCSTn: Neu5Acα2-6GalNAcα-O-Ser/Thr;
Core 2: GlcNAcβ1-6(Galβ1-3)GalNAc-α-O-Ser/Thr;
MUCTGlcNAc: GlcNAcβ1-3Galβ1-3GalNAcα-O-Ser/Thr,
MUCST: Neu5Acα2-3Galβ1-3GalNAc-α-O-Ser/Thr,
MUC6ST: Galβ1-3(Neu5Acα2-6)GalNAcα-O-Ser/Thr,
MUCdST: Neu5Acα2-3Galβ1-3(Neu5Acα2-6)GalNAcα-O-Ser/Thr

## Data availability

The COPASI files used for illustrative purposes are available at (https://github.com/scientificomputing/CytoCopasi).

## REFERENCES

1. Al Masri A, Gendler SJ. 2005. Muc1 affects c-Src signaling in PyV MT-induced mammary tumorigenesis. Oncogene 24(38):5799–5808.

2. Baeuerle PA, Baltimore D. 1988. IκB: a specific inhibitor of the NF-κB transcription factor. Science 242(4878):540–546.

3. Bangarh R, Khatana C, Kaur S, Sharma A, Kaushal A, Siwal SS, Tuli HS, Dhama K, Thakur VK, Saini RV. 2023. Aberrant protein glycosylation: Implications on diagnosis and Immunotherapy. Biotechnology Advances 66:108149.

4. Beatson R, Graham R, Freile FG, Cozzetto D, Kannambath S, Pfeifer E, Woodman N, Owen J, Nuamah R, Mandel U. 2020. Cancer-associated hypersialylated MUC1 drives the differentiation of monocytes into macrophages with a pathogenic phenotype. bioRxiv:2020.05. 06.080713.

5. Beatson R, Tajadura-Ortega V, Achkova D, Picco G, Tsourouktsoglou T-D, Klausing S, Hillier M, Maher J, Noll T, Crocker PR. 2016. The mucin MUC1 modulates the tumor immunological microenvironment through engagement of the lectin Siglec-9. Nature immunology 17(11):1273–1281.

6. Boll EJ, Ayala-Lujan J, Szabady RL, Louissaint C, Smith RZ, Krogfelt KA, Nataro JP, Ruiz-Perez F, McCormick BA. 2017. Enteroaggregative Escherichia coli adherence fimbriae drive inflammatory cell recruitment via interactions with epithelial MUC1. MBio 8(3):e00717–17.

7. Bowles WHD, Gloster TM. 2021. Sialidase and Sialyltransferase Inhibitors: Targeting Pathogenicity and Disease. Front Mol Biosci 8:705133.

8. Burchell J, Poulsom R, Hanby A, Whitehouse C, Cooper L, Clausen H, Miles D, Taylor-Papadimitriou J. 1999. An α2,3 sialyltransferase (ST3Gal I) is elevated in primary breast carcinomas. Glycobiology 9(12):1307–1311.

9. Chang K-H, Lee L, Chen J, Li W-S. 2006. Lithocholic acid analogues, new and potent α-2, 3- sialyltransferase inhibitors. Chemical communications (6):629–631.

10. Chen J-Y, Tang Y-A, Huang S-M, Juan H-F, Wu L-W, Sun Y-C, Wang S-C, Wu K-W, Balraj G, Chang T-T. 2011. A novel sialyltransferase inhibitor suppresses FAK/paxillin signaling and cancer angiogenesis and metastasis pathways. Cancer research 71(2):473–483.

11. Crous W, Naidoo KJ. 2016. Conformational and electrostatic analysis of SN1 donor analogue glycomimetic inhibitors of ST3Gal-I mammalian sialyltransferase. Bioorg Med Chem 24(20):4998–5005.

12. Deng J, Jiang R, Meng E, Wu H. 2022. CXCL5: A coachman to drive cancer progression. Frontiers in Oncology 12:944494.

13. Dziadek S, Griesinger C, Kunz H, Reinscheid UM. 2006. Synthesis and Structural Model of an α (2, 6)-Sialyl-T Glycosylated MUC1 Eicosapeptide under Physiological Conditions. Chemistry–A European Journal 12(19):4981–4993.

14. Figueiredo AS, Höfer T, Klotz C, Sers C, Hartmann S, Lucius R, Hammerstein P. 2009. Modelling and simulating interleukin-10 production and regulation by macrophages after stimulation with an immunomodulator of parasitic nematodes. The FEBS journal 276(13):3454–3469.

15. Gloster TM, Vocadlo DJ. 2012. Developing inhibitors of glycan processing enzymes as tools for enabling glycobiology. Nature Chemical Biology 8(8):683–694.

16. Hoops S, Sahle S, Gauges R, Lee C, Pahle J, Simus N, Singhal M, Xu L, Mendes P, Kummer U. 2006. COPASI—a complex pathway simulator. Bioinformatics 22(24):3067–3074.

17. Hsu C-C, Lin T-W, Chang W-W, Wu C-Y, Lo W-H, Wang P-H, Tsai Y-C. 2005. Soyasaponin-I-modified invasive behavior of cancer by changing cell surface sialic acids. Gynecologic oncology 96(2):415–422.

18. Hsu Y, Hou M, Kuo P, Huang Y, Tsai E. 2013. Breast tumor-associated osteoblast-derived CXCL5 increases cancer progression by ERK/MSK1/Elk-1/snail signaling pathway. Oncogene 32(37):4436–4447.

19. Kafi MA, Aktar K, Todo M, Dahiya R. 2020. Engineered chitosan for improved 3D tissue growth through Paxillin-FAK-ERK activation. Regenerative biomaterials 7(2):141–151.

20. Kawakami K, Scheidereit C, Roeder RG. 1988. Identification and purification of a human immunoglobulin-enhancer-binding protein (NF-kappa B) that activates transcription from a human immunodeficiency virus type 1 promoter in vitro. Proceedings of the National Academy of Sciences 85(13):4700–4704.

21. Kaya HE, Naidoo KJ. 2023. CytoCopasi: a chemical systems biology target and drug discovery visual data analytics platform. Bioinformatics 39(12):btad745.

22. Kennedy J, Eberhart R. Particle swarm optimization. Proceedings of ICNN’95-international conference on neural networks; 1995: ieee. p. 1942–1948.

23. Kouka T, Akase S, Sogabe I, Jin C, Karlsson NG, Aoki-Kinoshita KF. 2022. Computational modeling of O-linked glycan biosynthesis in CHO cells. Molecules 27(6):1766.

24. Krambeck FJ, Betenbaugh MJ. 2005. A mathematical model of N-linked glycosylation. Biotechnology and Bioengineering 92(6):711–728.

25. Li X, Wang M, Gong T, Lei X, Hu T, Tian M, Ding F, Ma F, Chen H, Liu Z. 2020. A S100A14- CCL2/CXCL5 signaling axis drives breast cancer metastasis. Theranostics 10(13):5687.

26. Lindén SK, Sheng YH, Every AL, Miles KM, Skoog EC, Florin TH, Sutton P, McGuckin MA. 2009. MUC1 limits Helicobacter pylori infection both by steric hindrance and by acting as a releasable decoy. PLoS pathogens 5(10):e1000617.

27. Liu G, Marathe DD, Matta KL, Neelamegham S. 2008. Systems-level modeling of cellular glycosylation reaction networks: O-linked glycan formation on natural selectin ligands. Bioinformatics 24(23):2740–2747.

28. Liu Z-X, Yu CF, Nickel C, Thomas S, Cantley LG. 2002. Hepatocyte growth factor induces ERK-dependent paxillin phosphorylation and regulates paxillin-focal adhesion kinase association. Journal of Biological Chemistry 277(12):10452–10458.

29. Lu Y, Liu J, Jia Y, Yang Y, Chen Q, Sun L, Song S, Huang L, Wang Z. 2019. Mass spectrometry analysis of changes in human milk N/O-glycopatterns at different lactation stages. Journal of agricultural and food chemistry 67(38):10702–10712.

30. Luo M, Hu Z, Kong Y, Li L. 2021. MicroRNA[z]432[z]5p inhibits cell migration and invasion by targeting CXCL5 in colorectal cancer. Experimental and therapeutic medicine 21(4):1–1.

31. Luo Y, Cao H, Lei C, Liu J. 2023. ST6GALNAC1 promotes the invasion and migration of breast cancer cells via the EMT pathway. Genes & Genomics 45(11):1367–1376.

32. Meany DL, Chan DW. 2011. Aberrant glycosylation associated with enzymes as cancer biomarkers. Clinical proteomics 8:1–14.

33. Monti P, Leone BE, Zerbi A, Balzano G, Cainarca S, Sordi V, Pontillo M, Mercalli A, Di Carlo V, Allavena P. 2004. Tumor-derived MUC1 mucins interact with differentiating monocytes and induce IL-10highIL-12low regulatory dendritic cell. The Journal of Immunology 172(12):7341– 7349.

34. Napoletano C, Rughetti A, Agervig Tarp MP, Coleman J, Bennett EP, Picco G, Sale P, Denda-Nagai K, Irimura T, Mandel U. 2007. Tumor-associated Tn-MUC1 glycoform is internalized through the macrophage galactose-type C-type lectin and delivered to the HLA class I and II compartments in dendritic cells. Cancer research 67(17):8358–8367.

35. Napoletano C, Zizzari IG, Rughetti A, Rahimi H, Irimura T, Clausen H, Wandall HH, Belleudi F, Bellati F, Pierelli L. 2012. Targeting of macrophage galactose-type C-type lectin (MGL) induces DC signaling and activation. European journal of immunology 42(4):936–945.

36. Nashed A, Naidoo KJ. 2024. Universal Glycosyltransferase Continuous Assay for Uniform Kinetics and Inhibition Database Development and Mechanistic Studies Illustrated on ST3GAL1, C1GALT1, and FUT1. ACS omega 9(15):17518–17532.

37. Orton RJ, Adriaens ME, Gormand A, Sturm OE, Kolch W, Gilbert DR. 2009. Computational modelling of cancerous mutations in the EGFR/ERK signalling pathway. BMC systems biology 3:1–17.

38. Patani N, Jiang W, Mokbel K. 2008. Prognostic utility of glycosyltransferase expression in breast cancer. Cancer genomics & proteomics 5(6):333–340.

39. Peterson JA, Hamosh M, Scallan CD, Ceriani RL, Henderson TR, Mehta NR, Armand M, Hamosh P. 1998. Milk fat globule glycoproteins in human milk and in gastric aspirates of mother’s milk-fed preterm infants. Pediatric Research 44(4):499–506.

40. Petzold L. 1983. Automatic selection of methods for solving stiff and nonstiff systems of ordinary differential equations. SIAM journal on scientific and statistical computing 4(1):136– 148.

41. Picco G, Julien S, Brockhausen I, Beatson R, Antonopoulos A, Haslam S, Mandel U, Dell A, Pinder S, Taylor-Papadimitriou J. 2010. Over-expression of ST3Gal-I promotes mammary tumorigenesis. Glycobiology 20(10):1241–1250.

42. Pothukuchi P, Agliarulo I, Russo D, Rizzo R, Russo F, Parashuraman S. 2019. Translation of genome to glycome: role of the Golgi apparatus. FEBS Lett 593(17):2390–2411.

43. Pribic J, Brazill D. 2012. Paxillin phosphorylation and complexing with Erk and FAK are regulated by PLD activity in MDA-MB-231 cells. Cellular Signalling 24(8):1531–1540.

44. Rughetti A, Pellicciotta I, Biffoni M, BaLJckstroLJm M, Link T, Bennet EP, Clausen H, Noll T, Hansson GC, Burchell JM. 2005. Recombinant tumor-associated MUC1 glycoprotein impairs the differentiation and function of dendritic cells. The journal of Immunology 174(12):7764– 7772.

45. Schomburg I, Chang A, Schomburg D. 2002. BRENDA, enzyme data and metabolic information. Nucleic acids research 30(1):47–49.

46. Sehl ME, Shimada M, Landeros A, Lange K, Wicha MS. 2015. Modeling of cancer stem cell state transitions predicts therapeutic response. PloS one 10(9):e0135797.

47. Senapathi T, Bray S, Barnett CB, Gruning B, Naidoo KJ. 2019. Biomolecular Reaction and Interaction Dynamics Global Environment (BRIDGE). Bioinformatics 35(18):3508–3509.

48. Senapathi T, Suruzhon M, Barnett CB, Essex J, Naidoo KJ. 2020. BRIDGE: An Open Platform for Reproducible High-Throughput Free Energy Simulations. Journal of Chemical Information and Modeling 60(11):5290–5295.

49. Shannon P, Markiel A, Ozier O, Baliga NS, Wang JT, Ramage D, Amin N, Schwikowski B, Ideker T. 2003. Cytoscape: a software environment for integrated models of biomolecular interaction networks. Genome research 13(11):2498–2504.

50. Silver DL, Naora H, Liu J, Cheng W, Montell DJ. 2004. Activated signal transducer and activator of transcription (STAT) 3: localization in focal adhesions and function in ovarian cancer cell motility. Cancer research 64(10):3550–3558.

51. Sp N, Kang DY, Joung YH, Park JH, Kim WS, Lee HK, Song K-D, Park Y-M, Yang YM. 2017. Nobiletin inhibits angiogenesis by regulating Src/FAK/STAT3-mediated signaling through PXN in ER+ breast cancer cells. International Journal of Molecular Sciences 18(5):935.

52. Suh J, Payvandi F, Edelstein LC, Amenta PS, Zong WX, Gélinas C, Rabson AB. 2002. Mechanisms of constitutive NF-κB activation in human prostate cancer cells. The Prostate 52(3):183–200.

53. Szabo R, Dobie C, Montgomery AP, Steele H, Yu H, Skropeta D. 2024. Synthesis of α-Hydroxy-1, 2, 3-Triazole-linked Sialyltransferase Inhibitors and Evaluation of Selectivity Towards ST3GAL1, ST6GAL1 and ST8SIA2. ChemMedChem 19(16):e202400088.

54. Teranishi S, Kimura K, Nishida T. 2009. Role of formation of an ERK-FAK-paxillin complex in migration of human corneal epithelial cells during wound closure in vitro. Investigative ophthalmology & visual science 50(12):5646–5652.

55. Tomczak K, Czerwińska P, Wiznerowicz M. 2015. Review The Cancer Genome Atlas (TCGA): an immeasurable source of knowledge. Contemporary Oncology/Współczesna Onkologia 2015(1):68–77.

56. Umaña P, Bailey JE. 1997. A mathematical model of N-linked glycoform biosynthesis. Biotechnology and bioengineering 55(6):890–908.

57. Urban MB, Baeuerle PA. 1990. The 65-kD subunit of NF-kappa B is a receptor for I kappa B and a modulator of DNA-binding specificity. Genes & development 4(11):1975–1984.

58. Wei M, Wynn R, Hollis G, Liao B, Margulis A, Reid BG, Klabe R, Liu PC, Becker-Pasha M, Rupar M. 2007. High-throughput determination of mode of inhibition in lead identification and optimization. SLAS Discovery 12(2):220–228.

59. Wen L, Zhang X, Zhang J, Chen S, Ma Y, Hu J, Yue T, Wang J, Zhu J, Wu T. 2020. Paxillin knockdown suppresses metastasis and epithelial[z]mesenchymal transition in colorectal cancer via the ERK signalling pathway. Oncology reports 44(3):1105–1115.

60. Wu C-Y, Hsu C-C, Chen S-T, Tsai Y-C. 2001. Soyasaponin I, a potent and specific sialyltransferase inhibitor. Biochemical and biophysical research communications 284(2):466–469.

61. Xu F, Zhao H, Li J, Jiang H. 2021. Mucin-type sialyl-Tn antigen is associated with PD-L1 expression and predicts poor clinical prognosis in breast cancer. Gland Surgery 10(7):2159.

62. Zandi E, Rothwarf DM, Delhase M, Hayakawa M, Karin M. 1997. The IkB kinase complex (IKK) contains two kinase subunits, IKKa and IKKb, necessary for IkB phosphorylation and NF-kB activation. Cell 91(2):243–252.

63. Zhang T, Ma C, Zhang Z, Zhang H, Hu H. 2021. NF-κB signaling in inflammation and cancer. MedComm 2(4):618–653.

64. Zhao J, Ou B, Han D, Wang P, Zong Y, Zhu C, Liu D, Zheng M, Sun J, Feng H. 2017. Tumor-derived CXCL5 promotes human colorectal cancer metastasis through activation of the ERK/Elk-1/Snail and AKT/GSK3β/β-catenin pathways. Molecular cancer 16:1–15.

65. Zhao Q, Barclay M, Hilkens J, Guo X, Barrow H, Rhodes JM, Yu L-G. 2010. Interaction between circulating galectin-3 and cancer-associated MUC1 enhances tumour cell homotypic aggregation and prevents anoikis. Molecular cancer 9:1–12.

66. Zhao Q, Lee FS. 1999. Mitogen-activated protein kinase/ERK kinase kinases 2 and 3 activate nuclear factor-κB through IκB kinase-α and IκB kinase-β. Journal of Biological Chemistry 274(13):8355–8358.

