## Supplementary material for "Integrated tumour-immune cell response modelling of Luminal A breast cancer details malignant signalling and ST3Gal1 inhibitor-induced reversal": Details of model construction and Tables listing reactions parameters and formulas

**Integrated breast cancer tumour immune response systems model reveals malignant signalling in Luminal A and a potential mitigating effect of an ST3Gal1 Inhibitor**

Hikmet Emre Kaya [1,2] and Kevin J. Naidoo [1,2,*]

[1]Scientific Computing Research Unit Address, PD Hahn Building, University of Cape Town, Rondebosch 7701, [2]Department of Chemistry, PD Hahn Building, University of Cape Town, Rondebosch 7701.

*To whom correspondence should be addressed.

### 1- Construction of the Models

A

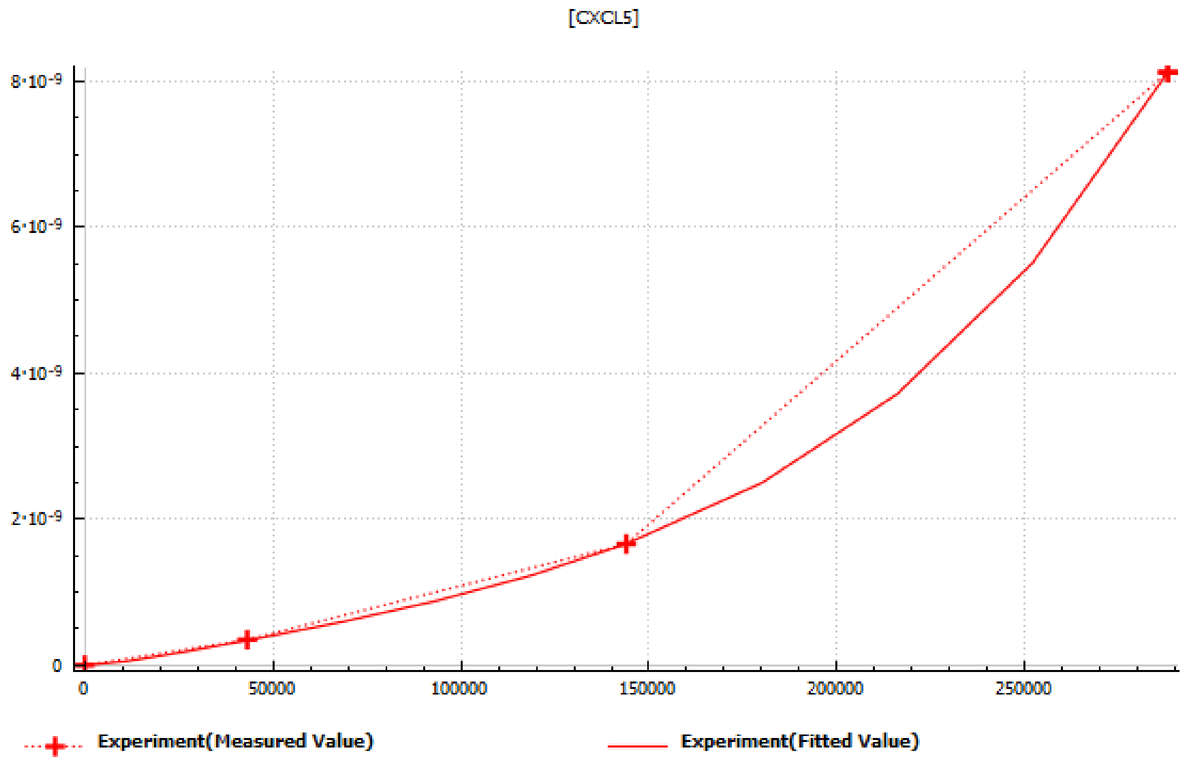

B

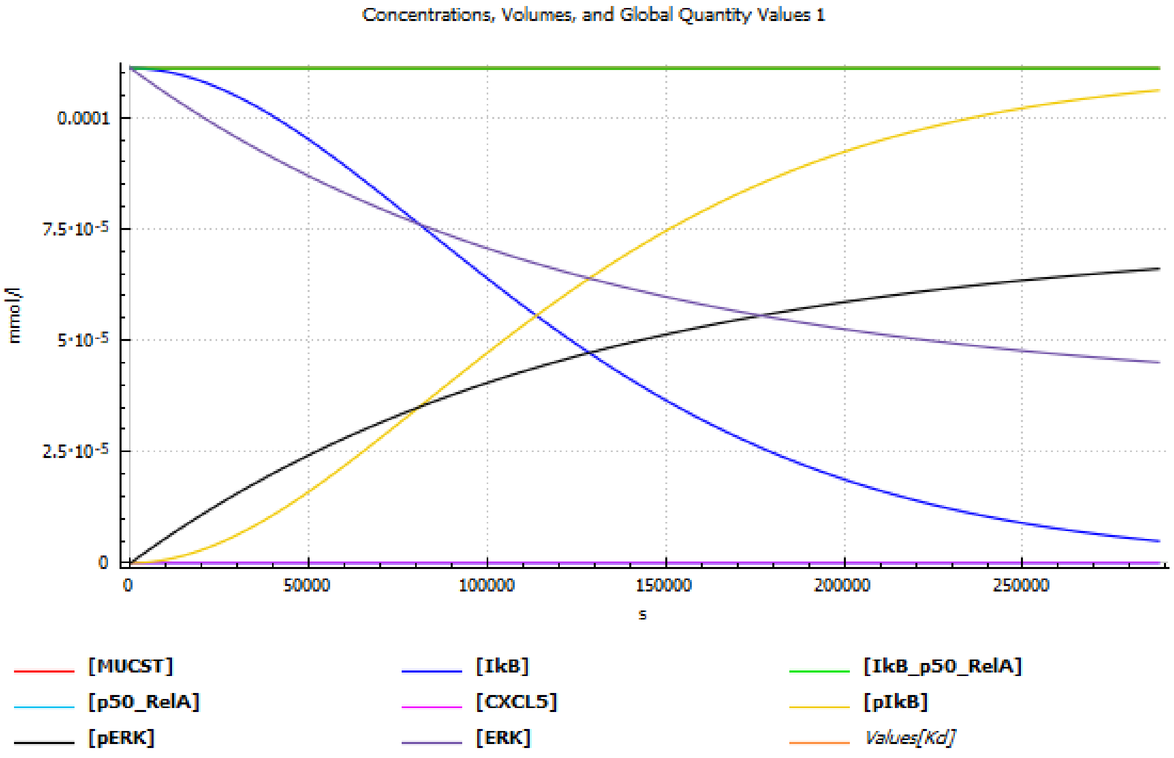

C

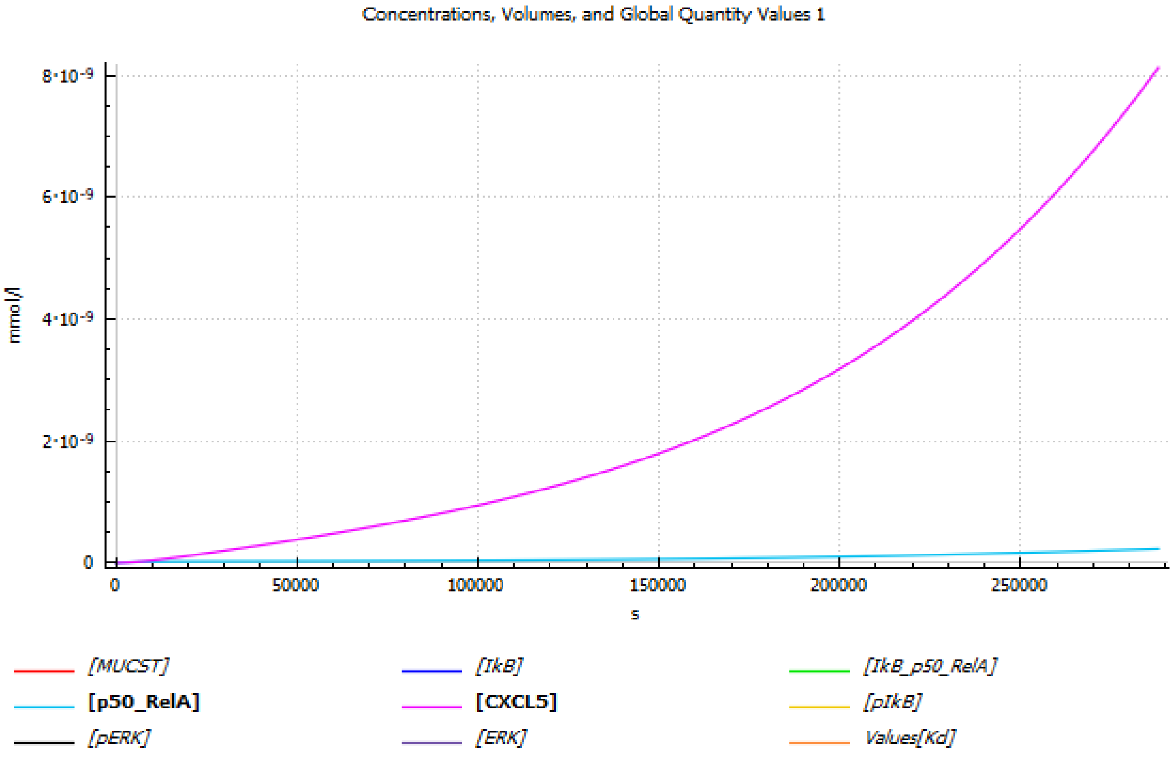

**Fig S1: The Time-Course Profiling of The Macrophage Network after Parameter Estimation.** (A) The fitted model exhibits an objective function of ~1 x 10 $^{-22}$, and fitted curve coincides with the experimentally determined values. (B) The time-course simulation of the parameter-estimated network illustrates the network shifting towards the phosphorylation of ERK and IkB, as their non-phosphorylated versions. (C) The time-course simulation depicts an exponential growth CXCL5 as p50_RelA gets activated.

**Table S1**

The reactions and parameters used in the construction of the glycosylation model

| Reaction/ Enzyme | Formula | $K_m$ (mM)[a] | $V_{max}$ (mM/s) [b] | Reference |
|---|---|---|---|---|
| POC1B-GALNT4 | MUC1 → MUCTn | 0.92 | $7.88 \times 10^{-5}$ | (Wandall et al. 1997) |
| C1GALT1 | MUCTn → MUCT | 0.00035 | 0.00074 | (Mendicino et al. 1982) |
| GCNT1 | MUCT → Core2 | 5.2 | $1.62 \times 10^{-5}$ | (Williams et al. 1980) |
| B3GNT3 | MUCT → MUCTGlcNAc | 1.2 | $2.38 \times 10^{-7}$ | (Schachter et al. 1989) |
| ST6GALNAC1 | MUCTn → MUCSTn | 2.93 | 0.00039 | (Sadler et al. 1979) |
| ST3GAL1 | MUCT → MUCST | 0.018 | $5.39 \times 10^{-6}$ | (Vallejo-Ruiz et al. 2001) |
| ST6GALNAC2_a | MUCT → MUC6ST | 2.93 | $5.39 \times 10^{-6}$ | (Sadler et al. 1979)[c] |
| ST6GALNAC2_b | MUCST → MUCdST | 2.93 | $1.13 \times 10^{-6}$ | (Sadler et al. 1979)[c] |

[a] Parameters were extracted from the BRaunschweig ENzyme Database (BRENDA) (Schomburg et al. 2002) using CytoCopasi (Kaya and Naidoo 2023).

[b] Parameters were estimated to based-on healthy breast milk metabolome (Lu et al. 2019)

[c] The Km value for ST6GALNAC1 was used due to the lack of acceptor-specific enzyme kinetics data for ST6GALNAC2

**Table S2**

The complete list of reactions, parameters, and initial concentration values of the monocyte-derived macrophage model

| Reaction | Chemical Formula | Rate Law | Parameters [a] |
|---|---|---|---|
| **R1** | MUCST + ERK → MUCST + pERK | k1 . [ERK] . [MUCST] | k1=0.0477239 l/(mmol.s) |
| **R2** | pERK → ERK | k2 . [pERK] | k2= $2.73077 \cdot 10^{-6}$ 1/s |
| **R3** | pERK + IkB → pERK + pIkB | k3 . [IkB] . [pERK] | k3 = 0.24184 l/(mmol.s) |
| **R4** | pIkB → IkB | k4 . [pIkB] | k4 = $5.65833 \cdot 10^{-12}$ 1/s |
| **R5** | IkB + p50_RelA → IkB_p50_RelA | k5 . [IkB] . [p50_RelA] | k5 = 1.28397 l/(mmol,s) |
| **R6** | IkB_p50_RelA → IkB + p50_RelA | k6 . [IkB_p50_RelA] | k6 = $3.08597 \cdot 10^{-11}$ 1/s |

| | | | |
|---|---|---|---|
| **R7** | p50_RelA → p50_RelA + CXCL5 | k7 . [p50_RelA] | k7 = 0.000354354 1/s |

[a] Parameters were fit to CXCL5 production in monocytes co-cultured with MUCST-bearing T-47D cell lines (Beatson et al. 2020)

**Table S3**

List of modifications to the reaction network proposed by Sehl et al. (Sehl et al. 2015) [a]

| **Reaction** | **Original Reaction** | **Modified Reaction** | **Parameter: $k_1$** |
|---|---|---|---|
| R1 | IL6 + gp130 → IL6_gp130 | CXCL5 + CXCR2 → CXCL5_CXCR2 | 0.0001 *1/(d. nM)* [a] |
| R3 | IL6_gp130 → IL6 + gp130 | CXCL5_CXCR2 → CXCL5 + CXCR2 | 0.008 *1/d* |
| R9 | IL6_gp130 + Stat3 → IL6_gp130 + aStat3 | CXCL5_CXCR2 + Stat3 → CXCL5_CXCR2 + aStat3 | 0.0001 *1/(d. nM)* |
| R19 | Lin28 + Let7 → Lin28_Let7 | Lin28 + mir432 → Lin28_mir432 | 0.0001 *1/(d. nM)* |
| R20 | Lin28_Let7 → Lin28 + Let7 | Lin28_mir432 → Lin28 + mir432 | 0.008 *1/d* |
| R25 | IL6_Let7 → IL6 + Let7 | mir432_CXCL5 → mir432+ CXCL5 | 0.008 *1/d* |
| R28 | IL6 + Let7 → IL6_Let7 | mir432+ CXCL5 → mir432_CXCL5 | 0.0001 *1/(d. nM)* |

**Table S4**

List of reactions added to the reaction network proposed by Sehl et al. (Sehl et al. 2015) [a]

| **Reaction Name** | **Reaction** | **Parameter: $k_1$** |
|---|---|---|
| R38 | CXCL5_CXCR2 + Erk → CXCL5_CXCR2 + aErk | 0.0001 *1/(d. nM)* [a] |
| R39 | aErk + IkB_p50_RelA → aErk + IkB + p50_RelA | 0.0001 *1/(d. nM)* [a] |
| R40 | CXCL5_CXCR2 + Akt → CXCL5_CXCR2 + aAkt | 0.0001 *1/(d. nM)* [a] |
| R41 | aErk → Erk | 0.008 *1/d* |

[a] The rate constants from the original study were used, but particle numbers were replaced with molar concentrations

### 2- Construction of the Luminal A-type MUC1 O-Glycosylation model via Perturbations

**Table S5**

The parameters of the Luminal A MUC1 O-Glycosylation Model based on The Cancer Genome Atlas (Tomczak et al. 2015) [a]

| Name | Perturbed Item | Fold Change | Perturbed Value |
|---|---|---|---|
| MUC1 | Initial Concentration [b] | 8.59 | 9.60 |
| B3GNT3 | $V_{max}$ [c] | 0.27 | $6.36 \times 10^{-8}$ |
| C1GALT1 | $V_{max}$ | 0.67 | $4.98 \times 10^{-4}$ |
| GALNT1 | $V_{max}$ | 1.36 | $1.07 \times 10^{-4}$ |
| ST3Gal1 | $V_{max}$ | 1.40 | $7.55 \times 10^{-6}$ |
| ST6GALNAC1 | $V_{max}$ | 0.27 | $1.05 \times 10^{-4}$ |
| ST6GALNAC2_a<br>ST6GALNAC2_b | $V_{max}$ | 1.25 | $7.40 \times 10^{-6}$<br>$1.41 \times 10^{-5}$ |

[a] The enzymes not shown in this table were classified as not significant (NS) in the TCGA dataset

[b] The unit is mM

[c] The unit is mM/s

### 3- Altered Concentration Profiles of the Networks in Luminal A compared to Normal Glycosylation

**Note: Please go to the CytoCopasi User Manual, section 8.4.3.1 if you wish to replicate the comparison results**

**Table S6**

Comparative Simulation of MUC1 O-Glycosylation: Normal vs LumA, t= 288000 s

| Metabolite | Normal (mM) | LumA (mM) | Change% (Normal vs LumA) |
|---|---|---|---|

| | | | |
|---|---|---|---|
| MUCTn | $2.69 \times 10^{-5}$ | $8.90 \times 10^{-5}$ | 230.41 ↑ |
| MUCT | 10.52 | 22.5 | 113.38 ↑ |
| MUCTGlcNAc | 0.0519 | 0.0155 | 70.08 ↓ |
| MUC6ST | 0.924 | 1.55 | 68.05 ↑ |
| MUCdST | 0.514 | 0.810 | 57.71 ↑ |
| Core2 | 2.20 | 2.91 | 32.36 ↑ |
| MUCST | 1.02 | 1.35 | 32.12 ↑ |
| MUCSTn | 0.00103 | $9.20 \times 10^{-4}$ | 10.95 ↓ |

**Table S7**

Impact of MUCST Upregulation on Monocyte Metabolites, including CXCL5: Normal vs LumA, t = 288000 s

| Metabolite | Before Perturbation (mM) | After Perturbation (mM) | Change% (Normal vs LumA) |
|---|---|---|---|
| IkB | $4.93 \times 10^{-6}$ | $2.75 \times 10^{-6}$ | 44.20 ↓ |
| MUCST | $1.11 \times 10^{-4}$ | $1.47 \times 10^{-4}$ | 32.12 ↑ |
| p50_RelA | $2.31 \times 10^{-10}$ | $2.92 \times 10^{-10}$ | 26.72 ↑ |
| CXCL5 | $8.12 \times 10^{-9}$ | $9.98 \times 10^{-9}$ | 22.84 ↑ |
| ERK | $4.50 \times 10^{-5}$ | $3.60 \times 10^{-5}$ | 20.02 ↓ |
| pERK | $6.60 \times 10^{-5}$ | $7.50 \times 10^{-5}$ | 13.66 ↑ |
| pIkB | $1.06 \times 10^{-4}$ | $1.08 \times 10^{-4}$ | 2.06 ↑ |
| IkB_p50_RelA | $1.11 \times 10^{-4}$ | $1.11 \times 10^{-4}$ | $5.55 \times 10^{-5}$ ↓ |

**Table S8**

Impact of CXCL5 Upregulation caused by MUCST-Siglec9 Binding on Intracellular Signaling: Normal vs LumA t= 3,33 days

| Metabolite | Before Perturbation (nM) | After Perturbation (nM) | Change% (Normal vs LumA) |
|---|---|---|---|
| aErk | 0.053 | 0.065 | 22.337 ↑ |
| mir605_CXCL5 | 3.171 | 3.872 | 22.103 ↑ |
| CXCL5_CXCR2 | 3.171 | 3.872 | 22.103 ↑ |
| aStat3 | 0.059 | 0.070 | 20.302 ↑ |
| aAkt | 0.112 | 0.123 | 10.633 ↑ |
| CXCR2 | 96.829 | 96.128 | 0.724 ↓ |
| mir605 | 96.787 | 96.087 | 0.724 ↓ |
| Lin28_mir605 | 0.042 | 0.041 | 0.494 ↓ |
| Erk | 99.946 | 99.934 | 0.0119 ↓ |
| Stat3 | 99.941 | 99.929 | 0.0119 ↓ |

| | | | |
|---|---|---|---|
| Akt | 99.888 | 99.876 | 0.0119 ↓ |
| Lin28 | 2.536 | 2.537 | 0.0080 ↑ |
| Lin28_HER2mRNA | 0.043 | 0.043 | 0.0040 ↑ |
| aBCatenin | 0.096 | 0.096 | 0.00180 ↑ |
| IkB_p50_RelA | 3.223 | 3.223 | $3.04 \times 10^{-4}$ ↓ |
| IkB | 96.777 | 96.777 | $1.01 \times 10^{-5}$ ↑ |
| p50_RelA | 96.777 | 96.777 | $1.01 \times 10^{-5}$ ↑ |
| HER2mRNA | 99.957 | 99.957 | $1.72 \times 10^{-6}$ ↓ |
| BCatenin | 99.904 | 99.904 | $1.72 \times 10^{-6}$ ↓ |
| HER2 | 96.830 | 96.830 | $9.62 \times 10^{-9}$ ↑ |
| IL8_CXCR1 | 3.181 | 3.181 | $1.39 \times 10^{-9}$ ↓ |
| HER2_EGFR | 3.170 | 3.170 | $2.55 \times 10^{-10}$ ↓ |
| CXCR1 | 96.819 | 96.819 | $4.56 \times 10^{-11}$ ↑ |
| IL8 | 96.819 | 96.819 | $4.56 \times 10^{-11}$ ↑ |
| EGFR | 96.830 | 96.830 | $8.28 \times 10^{-12}$ ↑ |

### 4- Impact of ST3Gal1 Inhibition by Soyasaponin-I on The Concentration Profiles

**Note: Please go to the CytoCopasi User Manual, section 8.4.3.2 if you wish to replicate the perturbation results**

**Table S9**

Impact of Soyasaponin-I inhibition on the Luminal A-type MUC1 O-Glycosylation Network

| Metabolite | Before Perturbation (mM) | After Perturbation (mM) | Change% (LumA vs Drug-treated Lum A) |
|---|---|---|---|
| MUCST | 1.35 | 0.993 | 26.59 ↓ |
| MUCdST | 0.810 | 0.643 | 20.70 ↓ |
| MUCSTn | $9.20 \times 10^{-4}$ | $9.52 \times 10^{-4}$ | 3.43 ↑ |
| MUCT | 22.5 | 23.0 | 2.23 ↑ |
| Core2 | 2.91 | 2.93 | 0.66 ↑ |
| MUC6ST | 0.155 | 0.156 | 0.46 ↑ |
| MUCTGlcNAc | 0.0155 | 0.0156 | 0.62 ↑ |

**Table S10**

The Downstream Impact of Soyasaponin-I inhibition on the Luminal A-type Macrophage Network

| **Metabolite** | **Before Perturbation (mM)** | **After Perturbation (mM)** | **Change% (Lum A vs Drug-treated Lum A)** |
|---|---|---|---|
| IkB | $4.93 \times 10^{-6}$ | $2.91 \times 10^{-5}$ | 82.22 ↑ |
| MUCST | $1.11 \times 10^{-4}$ | $8.15 \times 10^{-5}$ | 26.59 ↓ |
| p50_RelA | $2.31 \times 10^{-10}$ | $1.71 \times 10^{-10}$ | 25.87 ↓ |
| ERK | $4.50 \times 10^{-5}$ | $5.55 \times 10^{-5}$ | 23.26 ↑ |
| CXCL5 | $8.12 \times 10^{-9}$ | $6.46 \times 10^{-9}$ | 20.50 ↓ |
| pERK | $6.60 \times 10^{-5}$ | $5.55 \times 10^{-5}$ | 15.87 ↓ |
| pIkB | $1.06 \times 10^{-4}$ | $1.02 \times 10^{-4}$ | 3.82 ↓ |
| IkB_p50_RelA | $1.11 \times 10^{-4}$ | $1.11 \times 10^{-4}$ | $1.37 \times 10^{-4}$ ↑ |

**Table S11**

The Downstream Impact of Soyasaponin-I inhibition on the Luminal A-type Intracellular Signaling Network

| **Metabolite** | **Before Perturbation (nM)** | **After Perturbation (nM)** | **Change% (Lum A vs Drug-treated Lum A)** |
|---|---|---|---|
| aErk | 0.0532 | 0.0424 | 20.14 ↓ |
| mir605_CXCL5 | 3.171 | 2.538 | 19.97 ↓ |
| CXCL5_CXCR2 | 3.171 | 2.538 | 19.97 ↓ |
| CXCL5 | 96.278 | 77.05 | 19.97 ↓ |
| aStat3 | 0.0585 | 0.0478 | 18.31 ↓ |
| aAkt | 0.112 | 0.101 | 9.59 ↓ |
| CXCR2 | 96.82869 | 98.407 | 1.63 ↑ |
| mir605 | 96.78752 | 98.36468 | 1.63 ↑ |
| Lin28_mir605 | 0.041625 | 0.042076 | 1.08 ↑ |
| Erk | 99.94683 | 99.97347 | 0.0267 ↑ |
| Stat3 | 99.9415 | 99.96814 | 0.0267 ↑ |
| Akt | 99.88837 | 99.915 | 0.0267 ↑ |
| Lin28 | 2.536446 | 2.535999 | 0.0176 ↓ |
| Lin28_HER2mRNA | 0.042534 | 0.04253 | 0.0089 ↓ |
| aBCatenin | 0.095673 | 0.095669 | 0.0040 ↓ |
| IkB_p50_RelA | 3.222625 | 3.222646 | $6.68 \times 10^{-4}$ ↑ |

| | | | |
|---|---|---|---|
| IkB | 96.77738 | 96.77735 | 2.23 x $10^{-5}$ ↓ |
| p50_RelA | 96.77738 | 96.77735 | 2.23 x $10^{-5}$ ↓ |
| BCatenin | 99.90433 | 99.90433 | 3.79 x $10^{-6}$ ↑ |
| HER2mRNA | 99.95747 | 99.95747 | 3.79 x $10^{-6}$ ↑ |
| HER2 | 96.83019 | 96.83019 | 2.10 x $10^{-8}$ ↓ |
| HER2_EGFR | 3.170188 | 3.170188 | 4.16 x $10^{-9}$ ↓ |
| IL8_CXCR1 | 3.180592 | 3.180592 | 6.34 x $10^{-10}$ ↓ |
| EGFR | 96.82981 | 96.82981 | 1.36 x $10^{-10}$ ↑ |
| CXCR1 | 96.81941 | 96.81941 | 2.08 x $10^{-11}$ ↑ |
| IL8 | 96.81941 | 96.81941 | 2.08 x $10^{-11}$ ↑ |

### 5- Impact of ST3GAL1 Inhibition via Soyasaponin-I on Restoring the Normal State

**Note: Please go to the CytoCopasi User Manual, section 8.4.3.3 if you wish to replicate the comparison results**

**Table S12**

Comparison of the Concentration Profiles of Soyasaponin-I-treated Luminal A-type glycosylation to Normal Glycosylation

| Metabolite | Normal (mM) | Drug-treated Lum A (mM) | Change% (Normal vs Drug-treated Lum A) |
|---|---|---|---|
| MUCTn | 2.69 x $10^{-5}$ | 9.21 x $10^{-5}$ | 241.75 ↑ |
| MUCT | 10.52 | 22.96 | 118.14 ↑ |
| MUCTGlcNAc | 0.0519 | 0.0156 | 70.00 ↓ |
| MUC6ST | 0.924 | 1.556 | 68.83 ↑ |
| Core2 | 0.514 | 2.93 | 33.24 ↑ |
| MUCdST | 2.20 | 0.643 | 25.06 ↑ |
| MUCSTn | 1.02 | 9.52 x $10^{-4}$ | 7.89 ↓ |
| MUCST | 0.00103 | 0.993 | 3.01 ↓ |

**Table S13**

Comparison between the Concentration Profiles of the Macrophage Networks in Soyasaponin-I treated and normal states

| Metabolite | Normal (mM) | Drug-treated Lum A (mM) | Change% (Normal vs Drug-treated Lum A) |
|---|---|---|---|
| IkB | $4.93 \times 10^{-6}$ | $5.25 \times 10^{-6}$ | 6.40 ↑ |
| MUCST | $1.11 \times 10^{-4}$ | $1.08 \times 10^{-4}$ | 3.01 ↓ |
| p50_RelA | $2.31 \times 10^{-10}$ | $2.24 \times 10^{-10}$ | 2.78 ↓ |
| ERK | $4.50 \times 10^{-5}$ | $4.60 \times 10^{-5}$ | 2.28 ↑ |
| CXCL5 | $8.12 \times 10^{-9}$ | $7.94 \times 10^{-9}$ | 2.27 ↓ |
| pERK | $6.60 \times 10^{-5}$ | $6.50 \times 10^{-5}$ | 1.56 ↓ |
| pIkB | $1.06 \times 10^{-4}$ | $1.06 \times 10^{-4}$ | 0.30 ↓ |
| IkB_p50_RelA | $1.11 \times 10^{-4}$ | $1.11 \times 10^{-4}$ | $5.76 \times 10^{-6}$ ↑ |

**Table S14**

Comparison Between the Concentration Profiles of the Intracellular Signaling Networks in Soyasaponin-I-treated and Normal States

| Metabolite | Before Perturbation (nM) | After Perturbation (nM) | Change% (Normal vs Drug-treated Lum A) |
|---|---|---|---|
| aErk | 0.053 | 0.052 | 2.22 ↓ |
| CXCL5 | 96.278 | 94.153 | 2.21 ↓ |
| mir605_CXCL5 | 3.171 | 3.101 | 2.20 ↓ |
| CXCL5_CXCR2 | 3.171 | 3.102 | 2.20 ↓ |
| aStat3 | 0.059 | 0.057 | 2.02 ↓ |
| aAkt | 0.112 | 0.110 | 1.06 ↓ |
| CXCR2 | 96.829 | 96.899 | 0.0721 ↑ |
| mir605 | 96.788 | 96.857 | 0.0721 ↑ |
| Lin28_mir605 | 0.04162 | 0.04164 | 0.0480 ↑ |
| Erk | 99.947 | 99.948 | 0.00118 ↑ |
| Stat3 | 99.942 | 99.943 | 0.00118 ↑ |
| Akt | 99.888 | 99.89 | 0.00118 ↑ |
| Lin28 | 2.536446 | 2.536426 | $7.81 \times 10^{-4}$ ↓ |
| Lin28_HER2mRNA | 0.042534 | 0.042534 | $3.95 \times 10^{-4}$ ↓ |
| aBCatenin | 0.0957 | 0.0957 | $1.75 \times 10^{-4}$ ↓ |
| IkB_p50_RelA | 3.223 | 3.223 | $2.97 \times 10^{-5}$ ↑ |
| IkB | 96.777 | 96.777 | $9.88 \times 10^{-7}$ ↓ |
| p50_RelA | 96.777 | 96.777 | $9.88 \times 10^{-7}$ ↓ |
| BCatenin | 99.904 | 99.904 | $1.68 \times 10^{-7}$ ↑ |

| | | | |
|---|---|---|---|
| HER2mRNA | 99.957 | 99.957 | $1.68 \times 10^{-7}$ ↑ |
| HER2 | 96.830 | 96.830 | $9.32 \times 10^{-10}$ ↓ |
| HER2_EGFR | 3.170 | 3.170 | $2.97 \times 10^{-10}$ ↓ |
| IL8_CXCR1 | 3.181 | 3.181 | $1.30 \times 10^{-10}$ ↓ |
| EGFR | 96.830 | 96.830 | $9.69 \times 10^{-12}$ ↑ |
| CXCR1 | 96.819 | 96.819 | $4.27 \times 10^{-12}$ ↑ |
| IL8 | 96.819 | 96.819 | $4.27 \times 10^{-12}$ ↑ |

Beatson R, Graham R, Freile FG, Cozzetto D, Kannambath S, Pfeifer E, Woodman N, Owen J, Nuamah R, Mandel U. 2020. Cancer-associated hypersialylated MUC1 drives the differentiation of monocytes into macrophages with a pathogenic phenotype. *bioRxiv*:2020.05. 06.080713.

Kaya HE, Naidoo KJ. 2023. CytoCopasi: a chemical systems biology target and drug discovery visual data analytics platform. *Bioinformatics* 39(12):btad745.

Lu Y, Liu J, Jia Y, Yang Y, Chen Q, Sun L, Song S, Huang L, Wang Z. 2019. Mass spectrometry analysis of changes in human milk N/O-glycopatterns at different lactation stages. *Journal of agricultural and food chemistry* 67(38):10702-10712.

Mendicino J, Sivakami S, Davila M, Chandrasekaran E. 1982. Purification and properties of UDP-gal: N-acetylgalactosaminide mucin: beta 1, 3-galactosyltransferase from swine trachea mucosa. *Journal of Biological Chemistry* 257(7):3987-3994.

Sadler JE, Rearick JI, Hill RL. 1979. Purification to homogeneity and enzymatic characterization of an alpha-N-acetylgalactosaminide alpha 2 leads to 6 sialyltransferase from porcine submaxillary glands. *Journal of Biological Chemistry* 254(13):5934-5941.

Schachter H, Brockhausen I, Hull E. 1989. [30] High-performance liquid chromatography assays for N-acetylglucosaminyltransferases involved in N-and O-glycan synthesis. Methods in enzymology. Elsevier. p. 351-397.

Schomburg I, Chang A, Schomburg D. 2002. BRENDA, enzyme data and metabolic information. *Nucleic acids research* 30(1):47-49.

Sehl ME, Shimada M, Landeros A, Lange K, Wicha MS. 2015. Modeling of cancer stem cell state transitions predicts therapeutic response. *PloS one* 10(9):e0135797.

Tomczak K, Czerwińska P, Wiznerowicz M. 2015. Review The Cancer Genome Atlas (TCGA): an immeasurable source of knowledge. *Contemporary Oncology/Współczesna Onkologia* 2015(1):68-77.

Vallejo-Ruiz V, Haque R, Mir A-M, Schwientek T, Mandel U, Cacan R, Delannoy P, Harduin-Lepers A. 2001. Delineation of the minimal catalytic domain of human Galβ1-3GalNAc α2, 3-sialyltransferase (hST3Gal I). *Biochimica et Biophysica Acta (BBA)-Protein Structure and Molecular Enzymology* 1549(2):161-173.

Wandall HH, Hassan H, Mirgorodskaya E, Kristensen AK, Roepstorff P, Bennett EP, Nielsen PA, Hollingsworth MA, Burchell J, Taylor-Papadimitriou J. 1997. Substrate specificities of three members of the human UDP-N-acetyl-α-D-galactosamine: polypeptide N-acetylgalactosaminyltransferase family, GalNAc-T1,-T2, and-T3. *Journal of Biological Chemistry* 272(38):23503-23514.

Williams D, Longmore G, Matta K, Schachter H. 1980. Mucin synthesis. II. Substrate specificity and product identification studies on canine submaxillary gland UDP-GlcNAc: Gal beta 1-3GalNAc (GlcNAc leads to GalNAc) beta 6-N-acetylglucosaminyltransferase. *Journal of Biological Chemistry* 255(23):11253-11261.
